# Longitudinally stable, brain-based predictive models mediate the relationships between childhood cognition and socio-demographic, psychological and genetic factors

**DOI:** 10.1101/2021.02.21.432130

**Authors:** Narun Pat, Yue Wang, Richard Anney, Lucy Riglin, Anita Thapar, Argyris Stringaris

## Abstract

Cognitive abilities are one of the major transdiagnostic domains in the National Institute of Mental Health’s Research Domain Criteria (RDoC). Following RDoC’s integrative approach, we aimed to develop brain-based predictive models for cognitive abilities that a) are developmentally stable over years during adolescence and b) account for the relationships between cognitive abilities and socio-demographic, psychological and genetic factors. For this, we leveraged the unique power of the large-scale, longitudinal data from the Adolescent Brain Cognitive Development (ABCD) study (n ∼11k) and combined MRI data across modalities (task-fMRI from three tasks, resting-state fMRI, structural MRI, DTI) using machine-learning. Our brain-based, predictive models for cognitive abilities were stable across two years during young adolescence and generalisable to different sites, partially predicting childhood cognition at around 20% of the variance. Moreover, our use of ‘opportunistic stacking’ allowed the model to handle missing values, reducing the exclusion from around 80% to around 5% of the data. We found fronto-parietal networks during a working-memory task to drive childhood-cognition prediction. The brain-based, predictive models significantly, albeit partially, accounted for variance in childhood cognition due to (1) key socio-demographic and psychological factors (proportion mediated=18.65% [17.29%-20.12%]) and (2) genetic variation, as reflected by the polygenic score of cognition (proportion mediated=15.6% [11%-20.7%]). Thus, our brain-based predictive models for cognitive abilities facilitate the development of a robust, transdiagnostic research tool for cognition at the neural level in keeping with the RDoC’s integrative framework.

**Key Points:**

1. Using opportunistic stacking and multimodal MRI, we developed brain-based predictive models for children’s cognitive abilities that were longitudinally stable, generalisable to different sites and robust against missing data.
2. Our brain-based models were able to partially mediate the relationships of childhood cognitive abilities with the socio-demographic, psychological and genetic factors.
3. Our approach should pave the way for future researchers to employ multimodal MRI as a tool for the brain-based indicator of cognitive abilities, according to the integrative RDoC framework.

## Introduction

According to the Research Domain Criteria (RDoC), cognitive abilities are considered one of the major transdiagnostic domains, cutting across mental disorders^1^. In children and adults, cognitive abilities are related to various mental disorders, including but not limited to depression^2^, ADHD^3^ and psychotic disorders^4^. Cognitive abilities that span across cognitive tasks, such as language, mental flexibility, and memory, reflect a trait, known as general cognition or the *g*-factor^5^. Yet, we still don’t have predictive models that can robustly capture the relationship between *g*-factor and the brain. Having a brain-based predictive model for the *g*-factor is a key for us to adapt the RDoC’s integrative approach–to understand cognitive abilities across units of analyses, from behaviours to brain and genes that reflect the influences of socio-demographical and psychological factors across the lifespan^6, 7^.

Developing the brain-based predictive models for children’s *g*-factor to be used in the RDoC framework faces several challenges. The first challenge is longitudinal stability, which is one of the requirements in the RDoC framework^6, 7^. Predictive models should not only be generalisable to out-of-sample data (i.e., be predictive of children’s *g*-factor that were not part of the original sample) but also be developmentally stable^8^ in order to capture the *g*-factor across the lifespan^9^. Here we started to tackle this challenge by using--for the first time to our knowledge--longitudinal, large-scale data in children, from the Adolescent Brain Cognitive Development (ABCD) study^10^, to demonstrate the longitudinal stability of the brain-based predictive models across two years during adolescence.

The second challenge is multimodal integration. So far brain-based predictive models have been mainly built from a single MRI modality without integrating different sources of information from different MRI modalities. For instance, the *g*-factor is associated with activity during certain cognitive tasks, such as working memory^11, 12^(task-based functional MRI; task-fMRI), the intrinsic functional connectivity between different areas^13–15^(resting-state fMRI; rs-fMRI) and the anatomy of grey matter^16^ (structural MRI; sMRI) and white matter^17, 18^ (Diffusion Tensor Imaging; DTI). However, recent findings, mainly in adults, have started to show the benefits of integrating data across modalities, rather than relying solely on a single modality^8, 19, 20^. Here we adapted a machine-learning framework, called stacking^21^, to integrate information across MRI modalities into a ‘stacked’ model. Briefly, we separately built models to predict the *g*-factor based on each brain modality, resulting in one predicted value from each modality for each participant. We then built a ‘stacked’ model to predict the *g*-factor based on these predicted values. We tested if the stacked model indeed enhanced predictive performance over single modalities in predicting children’s *g*-factor.

The third challenge is missing data. Children’s neuroimaging data are notoriously affected by movement artifacts^22^. For example, the ABCD study recommended a set of quality control variables for detecting noisy data from each modality^10, 23^, resulting in a listwise exclusion of 17% to over 50% of data depending on a modality. If we were to exclude children who have noisy data from any single modality, we would have to exclude almost 80% of the data, strictly limiting the generalisability of our model to children with highly clean data (who are unlikely to be representative of the rest of the sample). We overcame this problem by using a recently developed framework, built on top of the stacking framework, called ‘opportunistic stacking’^24^. Briefly, we first duplicated predicted values from each modality-specific model, and imputed the missing value in each duplicate either with an arbitrarily high or low value. We then used Random Forest^25^ to create a final prediction from the imputed, predicted values. Accordingly, opportunistic stacking allows us to keep the data as long as there is at least one modality available, leaving more data in the model-building process and reducing the risk of missing-data bias.

Beyond demonstrating a robust out-of-sample relationship between the brain and the *g*-factor, the brain-based predictive models have to demonstrate the construct validity, especially for them to be used according to the RDoC framework^6^. For instance, RDoC stipulates that cognitive abilities are affected by socio-demographic and psychological factors^1, 26^. This is in line with recent studies showing that cognitive abilities are related to factors such as socio-economic status^27^, mental health^28, 29^ and extracurricular activities^30^. Accordingly, for the brain-based predictive models to demonstrate RDoC’s construct validity, the brain-based predictive models should be able to explain the associations between the *g*-factor and these socio-demographic and psychological factors.

Likewise, RDoC stipulates that cognitive abilities should not be studied as a unitary construct, but should rather be studied through different units of analysis, from behaviours to the brain and genes^1, 6, 7, 31^. Thus, the brain-based predictive models for cognitive abilities should be related to the “gene-based” predictive models for cognitive abilities, given that they both reflect different units of analysis of the same RDoC’s domain. A polygenic score (PGS), a composite measure of common gene variants, can be considered a predicted value from the gene-based predictive models^32^. For cognitive abilities^33^, a PGS is based on the associations between several Single Nucleotide Polymorphisms (SNPs) and cognitive abilities in a separate Genome-Wide Association Study (GWAS)^34^, such as in a recent GWAS among 257,841 adults^35^. Accordingly, for the brain-based predictive models to demonstrate RDoC’s construct validity, the brain-based predictive models should also be able to explain the associations between the *g*-factor and the PGS of cognitive abilities^35^.

To develop brain-based predictive models for the *g*-factor, we (i) used behavioural performance from cognitive tasks to derive a *g*-factor and (ii) built brain-based predictive models to predict this behaviourally derived *g*-factor from multimodal MRI data. We used the ABCD Release 3.0^10^, including baseline data (age 9-10 years old) from over 11,000 children and follow-up data (age 11-12 years old) from roughly half of the participants. We first derived children’s *g*-factor from their behavioural performance on six cognitive tasks using confirmatory factor analysis (CFA). We then built brain-based predictive models by treating multimodal MRI data as the features and the children’s *g*-factor derived from behavioural performance as the target. More specifically, in our models, we implemented opportunistic stacking^24^ to integrate MRI data across modalities and to deal with missing values from each modality. There were six modalities in total: three task-based fMRI (working-memory “N-Back”, reward “Monetary Incentive Delay; MID” and inhibitory control “Stop Signal”), rs-fMRI, sMRI, and DTI. To determine the robustness and longitudinal stability of the brain-based predictive models, we tested how well the models predicted the *g*-factor of unseen children at the same ages and at two years older as well as at different data-collection sites. Next, to demonstrate whether multimodal integration led to better predictive performance, we applied bootstrapping to compare the stacked model with the best-performing modality-specific model. To explain the feature importance of the final models (i.e., determining brain features that contributed highly to the prediction of the *g*-factor), we applied several ‘explainers’, including eNetXplorer^36^, conditional permutation importance^37^ and SHapley Additive exPlanations (SHAP)^38^.

We then conducted mediation analyses to ensure that the brain-based predictive models for the *g*-factor demonstrated RDoC’s construct validity. In these analyses, we tested the extent to which our brain-based predictive models could account for the relationships between the behaviourally derived *g*-factor and key socio-demographic, psychological and genetic factors. For this purpose, in addition to the brain-based predictive models, we also computed two additional predictive models that predicted the behaviourally derived *g*-factor, either from (a) 70 socio-demographic and psychological variables^30^ or (b) genes via a PGS of cognitive abilities^35^. The 70 socio-demographic and psychological variables covered children’s and/or their parents’ socio-demographics, mental health, personality, sleep, physical activity, screen use, drug use, developmental adversity and social interaction. This resulted in three predicted values of the *g*-factor, based on features of the predictive models: “brain-based *g*-factor”, “socio-demography-and-psychology-based *g*-factor” and “gene-based *g*-factor”. We then computed these predicted values on unseen children at each held-out data collection site and applied the mediation analyses. Here we treated (i) the socio-demography-and-psychology-based and gene-based *g*-factors as the independent variables, (ii) the brain-based *g*-factor as the mediator and (iii) the behaviourally derived *g*-factor as the dependent variable. Through these mediation analyses, we quantified the extent to which the brain-based predictive models developed in this study mediated the relationships between the behaviourally derived *g*-factor and socio-demographic, psychological and genetic factors.

## Methods

We employed the ABCD Study Curated Annual Release 3.0^10^, which included 3T MRI data and cognitive tests from 11,758 children (female=5,631) at the baseline (9-10 years old) and 5693 children (female = 2,617) at the two-year follow-up (11-12 years old). The study recruited the children from 21 sites across the United States^39^. We further excluded 54 children based on Snellen Vision Screener^40, 41^. These children either could not read any line, could only read the 1st (biggest) line, or could read up to the 4th line but indicated difficulty in reading stimuli on the iPad used for administering cognitive tasks (see below). The ethical considerations of the ABCD study, such as informed consent, confidentiality and communication with participants about assessment results, have been detailed elsewhere^42^. Institutional Review Boards where the data were collected approved the study’s protocols.

### The *g*-factor

We derived the *g*-factor using children’s behavioural performance from six cognitive tasks. These six tasks, collected on an iPad during a 70-min in-session visit outside of MRI^41, 43^, were available in both baseline and follow-up datasets. First, the Picture Vocabulary measured vocabulary comprehension and language^44^. Second, the Oral Reading Recognition measured reading and language decoding^45^. Third, the Flanker measured conflict monitoring and inhibitory control^46^. Fourth, the Pattern Comparison Processing measured the speed of processing^47^. Fifth, the Picture Sequence Memory measured episodic memory^48^. Sixth, the Rey-Auditory Verbal Learning measured memory recall after distraction and a short delay^49^.

Similar to the previous work^43, 50, 51^, we applied the 2^nd^-order model of the *g*-factor using confirmatory factor analysis (CFA) to encapsulate the *g*-factor as the higher-order latent variable underlying performance across cognitive tasks. More specifically, our input data were standardised performance from each cognitive task. In our 2^nd^-order model, we had the *g*-factor as the 2^nd^-order latent variable. We also had three 1^st^-order latent variables in the model: language (underlying the Picture Vocabulary and Oral Reading Recognition), mental flexibility (underlying the Flanker and Pattern Comparison Processing), and memory recall (underlying the Picture Sequence Memory and Rey-Auditory Verbal Learning).

We fixed latent factor variances to one and applied robust maximum likelihood estimation (MLR) with robust (Huber-White) standard errors and scaled test statistics. To demonstrate model fit, we used scaled and/or robust indices, including comparative fit index (CFI), Tucker-Lewis Index (TLI), root mean squared error of approximation (RMSEA) and Standardized Root Mean Square Residual (SRMR) as well as used internal consistency, OmegaL2^52^, of the *g*-factor. To implement the CFA, we used lavaan^53^ (version=.6-6) and semTools^52^ along with semPlot^54^ for visualisation. Note to ensure the robustness of the chosen *g*-factor model, we also examined the similarity in factor scores of the *g*-factor based on three different CFA models: the 2^nd^-order model, the single-factor model, and the mixture between Exploratory Factor Analysis (EFA) and CFA models (Supplementary Appendix 1).

### Multimodal MRI

We used MRI data from six modalities: three task-based fMRI, rs-fMRI, sMRI, and DTI. Note ‘modalities’ here referred to sets of features in our predictive models, as such we treated three task-based fMRI as separate modalities even though they were task-based fMRI. The ABCD Study provided detailed procedures on data acquisition and MRI image processing elsewhere^10, 23, 55^. We strictly followed their recommended exclusion criteria based on automated and manual QC review for each modality, listed under the *abcd_imgincl01* table^10^. The ABCD created an exclusion flag for each modality (with a prefix ‘imgincl’) based on several criteria, involving image quality, MR neurological screening, behavioural performance, number of Repetition Time (TRs) among others. We removed participants with an exclusion flag at any MRI indices, separately for each modality. We also applied the three interquartile range (3xIQR) rule (i.e., datapoint with a value over 3 IQRs away from the nearest quartile) with listwise deletion to remove observations with outliers in any indices within each modality. Additionally, to adjust for between-site variability, we used an Empirical Bayes method, ComBat^56, 57^. We applied ComBat to all modalities except for task-based fMRI, given that between-site variability was found to be negligible for task-based contrasts^57^. See below for our approach to mitigate data leakage due to 3xIQR and ComBat.

#### 1-3) Three Task-Based fMRI

We used task-based fMRI from three tasks. First, in the working-memory “N-Back” task^55, 58^, children saw pictures of houses and emotional faces. Depending on the blocks, children reported if a picture matched either: (a) a picture that is shown 2 trials earlier (2-back), or (b) a picture that is shown at the beginning of the block (0-back). To focus on working-memory-related activity, we use the [2-back vs 0-back] linear contrast (i.e., high vs. low working memory load). Second, in the Monetary Incentive Delay (MID) task^55, 59^, children needed to respond before the target disappeared. And doing so would provide them with a reward, if and only if the target followed the “reward cue” (but not in “neural cue”). To focus on reward anticipation-related activity, we used the [Reward Cue vs Neutral Cue] linear contrast. Third, in the Stop-Signal Task (SST)^55, 60^, children needed to withhold or interrupt their motor response to a “Go” stimulus when it is followed unpredictably by a Stop signal. To focus on inhibitory control-related activity, we used the [Any Stop vs Correct Go] linear contrast. Note that, for the SST, we used two additional exclusion criteria, *tfmri_sst_beh_glitchflag,* and *tfmri_sst_beh_violatorflag,* to address glitches in the task as recommended by the study^61, 62^. For all tasks, we used the average contrast values across two runs. More specifically, these contrasts were unthresholded, similar to previous work^63^. These values were embedded in the brain parcels based on FreeSurfer^64^’s Destrieux^65^ and ASEG^66^ atlases (148 cortical surface and 19 subcortical volumetric regions, resulting in 167 features for each task-based fMRI task).

#### 4) Resting-State fMRI (rs-fMRI)

During rs-fMRI collection, the children viewed a crosshair for 20 minutes. The ABCD’s preprocessing strategy has been published elsewhere^23^. Briefly, the study parcellated regions into 333 cortical-surface regions^67^ and correlated their time-series^23^. They then grouped these correlations based on 13 predefined large-scale networks^67^: auditory, cingulo-opercular, cingulo-parietal, default-mode, dorsal-attention, frontoparietal, none, retrosplenial-temporal, salience, sensorimotor-hand, sensorimotor-mouth, ventral-attention, and visual networks. Note that “none” refers to regions that do not belong to any networks. After applying Fisher r to z transformation, the study computed mean correlations between pairs of regions within each large-scale network (n=13) and between large-scale networks (n=78) and provided these mean correlations in their Releases^10^. This resulted in 91 features for the rs-fMRI. Given that the correlations between (not within) large-scale networks were highly collinear with each other (e.g., the correlation between auditory and cingulo-opercular was collinear with that between auditory and default-mode), we further decorrelated them using partial correlation. We first applied inverse Fisher r to z transformation, then partial correlation transformation, and then reapplied Fisher r to z transformation.

#### 5) Structural MRI (sMRI)

The ABCD study processed sMRI, including cortical reconstruction and subcortical volumetric segmentation, using FreeSurfer^64^. Here we considered FreeSurfer-derived Destrieux^65^ regional cortical thickness measures (n=148 cortical surface) and ASEG^66^ regional subcortical volume measures (n=19), resulting in 167 features for sMRI. We also adjusted regional cortical thickness and volumetric measures using mean cortical thickness and total intracranial volume, respectively.

#### 6) Diffusion Tensor Imaging (DTI)

Here we focused on fractional anisotropy (FA)^68^. FA characterises the directionality of the distribution of diffusion within white matter tracts, which can indicate the density of fibre packing^68^. The ABCD study segmented major white matter tracts using AtlasTrack^23, 69^. Here we considered FA of 23 major tracks, 10 of which were separately labelled for each hemisphere. These tracks included corpus callosum, forceps major, forceps minor, cingulate and parahippocampal portions of cingulum, fornix, inferior frontal occipital fasciculus, inferior longitudinal fasciculus, pyramidal/corticospinal tract, superior longitudinal fasciculus, temporal lobe portion of superior longitudinal fasciculus, anterior thalamic radiations and uncinate. This left 23 features for DTI.

### Predictive Models of Multimodal MRI: Opportunistic Stacking

To integrate multimodal MRI into one predictive model and to control for missing values across modalities, we applied opportunity stacking^24^ (Figure 1). We started with the first-layer training set. Here we used standardised features from each modality to separately predict the *g*-factor via a penalised regression. The main advantage of a penalised regression is its ease of interpretation given that the prediction is made based on a weighted sum of features. Moreover, predictive performance of penalised regressions for capturing brain-and-behaviour relationships in MRI appeared good, often on-par with other more black-box algorithms^15, 19, 24, 70, 71^. Following previous research^15^, we used Elastic Net^72^, a general form of penalised regression via the glmnet package^73^. Elastic Net requires two hyperparameters. First, the ‘penalty’ determines how strong the feature’s slopes are regularised. Second, the ‘mixture’ determines the degree to which the regularisation is applied to the sum of squared coefficients (known as Ridge) vs. to the sum of absolute values of the coefficients (known as LASSO). We tuned these two hyperparameters using a 10-fold cross-validation grid search and selected the model with the lowest Mean Absolute Error (MAE). In the grid, we used 200 levels of the penalty from 10^-10 to 10, equally spaced on the logarithmic-10 scale and 11 levels of the mixture from 0 to 1 on the linear scale.

**Figure 1.**
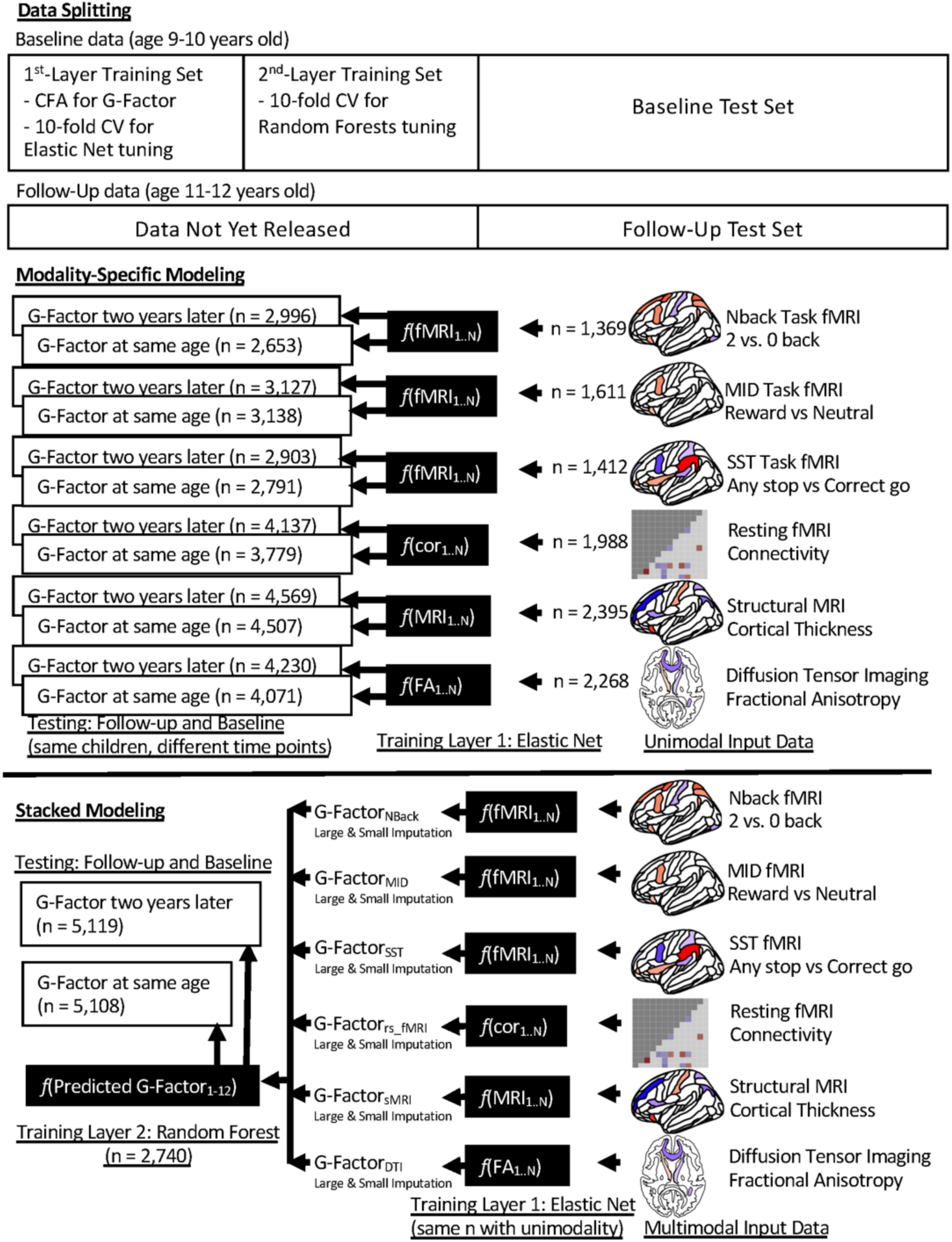
Longitudinal Predictive Modelling Approach Used for Out-of-sample Predictive Ability of Multimodal MRI. We split the data into four sets: first-layer training, second-layer training, baseline test, and follow-up test. We used the same participants in the baseline test and follow-up test sets. Modality-specific modelling only used the first-layer training set, while stacked modelling used both training sets to combine predicted values across modalities. At the first training layer, using Elastic Net, we separately predicted the g-factor based on each of the six modalities, resulting in six predicted values. At the second training layer, we applied opportunistic stacking by duplicating these six predicted values, and then imputed missing observations in one as an arbitrarily large value of 1000 and in the other as an arbitrarily small value of −1000, resulting in 12 predicted values. We then used Random Forest to predict the g-factor based on these 12 predicted values. The number of observations was different depending on the quality control of data from each modality. “Data not yet released” reflects the fact that ABCD Release 3.0^10^ only provided half of the follow-up data (age 11-12 years old), while providing the full baseline data (age 9-10 years old). CFA = Confirmatory Factor Analysis; CV = Cross-Validation; cor = correlation; FA = fractional anisotropy.

Once we obtained the final modality-specific models from the first-layer training set, we fit these models to data in the second-layer training set. This gave us six predicted values of the *g*-factor from six modalities, and these are the features to predict the *g*-factor in the second-layer training set. To handle missing observations when combining these modality-specific features, we applied the opportunistic stacking approach^24^ by creating duplicates of each modality-specific feature. After standardisation, we coded missing observations in one as an arbitrarily large value of 1000 and in the other as an arbitrarily small value of −1000, resulting in 12 features. That is, as long as a child had at least one modality available, we would be able to include this child in stacked modelling.

Previous research^24^ advocated for a more flexible algorithm that can capture non-linear and interactive relationships at the second-layer training set. Here we used the Random Forests algorithm^25^ from the ranger package^74^ to predict the *g*-factor from the 12 features^24, 75^.

Random Forests use a multitude of decision trees on various sub-samples of the data and implement averaging to enhance prediction and to control over-fitting. We used 1000 trees and turned two hyperparameters. First ‘mtry’ is the number of features randomly sampled at each split. Second ‘min_n’ is the minimum number of observations in a node needed for the node to be split further. We implemented a 10-fold cross-validation grid search and selected the model with the lowest Root Mean Squared Error (RMSE). In the grid, we used 12 levels of the mtry from 1 to 12, and 101 levels of the min_n from 1 to 1000, both on the linear scale. This resulted in the “stacked” model that incorporated data across modalities.

To prevent data leakage, we fit the CFA model to the observations in the first-layer training data and then computed factor scores of the *g*-factor on all training and test data. Note that to demonstrate the stability of the factor scores of the *g*-factor when applied to unseen data (i.e., not part of the modelling process), we also compared the factor scores of the *g*-factor estimated from the first-layer training data and the scores estimated from the whole baseline data (see Supplementary Appendix 2). Similarly, we also applied the 3xIQR rule and Combat separately for first-layer training, second-layer training, baseline test, and follow-up test data. For the machine learning workflow, we used ‘tidymodels’ (www.tidymodels.org).

### Testing the Robustness of the Predictive Models of Multimodal MRI

We examined the predictive ability of the models based on multimodal MRI between predicted vs. observed *g*-factor, using Pearson’s correlation (*r*), coefficient of determination (*R*^2^, calculated using the sum of square definition), mean absolute error (MAE), and root mean squared error (RMSE). To investigate the predictive ability of the modality-specific models, we used the models tuned from the first-layer training set. To investigate the predictive ability of the stacked model, we used the model tuned from both the first-layer and second-layer training sets.

#### 1) Out-of-sample Predictive Ability of Multimodal MRI: Baseline and Follow-Up Samples

We first split the data into four parts (Figure 1): 1) first-layer training set (n=3,041), 2) second-layer training set (n=3,042), 3) baseline test set (n=5,622) and 4) follow-up test set (n=5,656). Especially noteworthy is that children who were in the baseline test set were also in the follow-up test set. In other words, none of the children in the first-layer and second-layer training sets was in either of the test sets. We used the baseline test set for out-of-sample, same-age predictive ability, while we used the follow-up test sets for out-of-sample, longitudinal predictive ability.

To examine the performance of opportunistic stacking as a function of missing values, we further split the test sets based on the presence of each modality. First, Stacked All required data with at least one modality present. This allowed us to examine the stacked model’s performance when the missing values were all arbitrarily coded. Second, Stacked Complete required data with all modalities present. This represents the situation when the data were as clean as possible. Third, Stacked Best had the same missing values as the modality with the best prediction. This allowed us to make a fair comparison in performance between the stacked model and the model with the best modality, given their same noise level from missing value. Fourth, Stacked No Best did not have any data from the modality with the best prediction and had at least one modality present. This represents the highest level of noise possible.

#### 2) Comparing Out-of-sample Predictive Ability of Multimodal MRI between the Stacked Model and the Model with the best Modality: Baseline and Follow-Up Samples

Here we made a statistical comparison in the out-of-sample predictive ability between Stacked Best and the modality-specific model with the highest predictive performance, two of which had the same number of missing values in the test sets. We applied bootstrapping with 5,000 iterations to examine the differences in performance indices (including, *r*, *R*^2^, MAE, and RMSE) on both baseline and follow-up test sets. If stacking truly led to enhanced predictive performance, then we should see 95%CI of the bootstrapped differences to be different from 0.

#### 3) Out-of-site Predictive Ability of Multimodal MRI

To examine out-of-site predictive ability, we applied leave-one-site-out cross-validation to the baseline data. This enabled us to extract predicted values of the *g*-factor based on multimodal MRI data at each held-out site, and in turn, to examine the generalisability of different models on different data collection sites. Different sites involved different MRI machines, experimenters as well as demographics across the US^39^. Moreover, using leave-one-site-out cross-validation also prevented having the participants from the same family in the training and test sets. Here, we first removed data from one site that only recruited 34 participants and removed participants from six families who were separately scanned at different sites. We then held out data from one site as a test set and divided the rest to be first- and second-layer training sets. We cross-validated predictive ability across these held-out sites. We applied the same modelling approach with the out-of-sample predictive models, except for two configurations to reduce the amount of ram used and computational time. Specifically, in our grid search, we used 100 levels of penalty (as opposed to 200) for Elastic Net and limited the maximal min_n to 500 (as opposed to 1000) for Random Forests. For the stacked model, we tested its predictive ability on children with at least one modality (i.e., stacked all). We examined the out-of-site prediction between predicted vs. observed *g*-factor at each held-out site.

### Feature Importance of Multimodal MRI Models

To understand which features contribute to the prediction of the modality-specific (i.e., Elastic Net) models, we applied permutation from the eNetXplorer^76^ package to the first-layer training set of the out-of-sample predictive ability splits (Figure 1). We first chose the best mixture from the previously run grid and fit two sets of several Elastic Net models. The first “target” models used the true *g*-factor as the target, while the second “null” models used the randomly permuted *g*-factor as the target. eNetXplorer split the data into 10 folds 100 times/runs. For each run, eNetXplorer performed cross-validation by repeatedly training the target models on 9 folds and tested on the leftover fold. Also, in each cross-validation run, eNetXplorer trained the null models 25 times. eNetXplorer then used the mean of non-zero model coefficients across all folds in a given run as a coefficient for each run, *k^r^*. Across runs, eNetXplorer weighted the mean of a model coefficient by the frequency of obtaining a non-zero model coefficient per run. Formally, we defined an empirical p-value as:

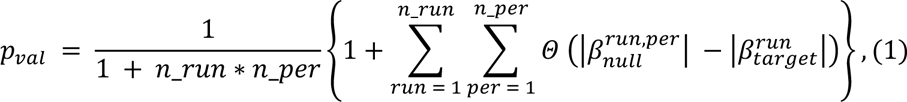

where *p_val_* is an empirical p-value, *run* is a run index, *n_run* is the number of runs, *per* is a permutation index, *n_per* is the number of permutation, Θ is the right-continuous Heaviside step function, and |β| is the magnitude of feature’ coefficient. That is, to establish statistical significance for each feature, we used the proportion of runs in which the null models performed better than the target models. We plotted the target models’ coefficients with *p_val_* < .05 on the brain images using the ggseg^77^ package.

To identify which modalities contributed strongly to the prediction of the stacked (i.e., Random Forests) model, we applied two methods: 1) conditional permutation importance (CPI)^78^ and 2) SHapley Additive exPlanations (SHAP)^38^ to the second-layer training set. CPI is an explainer, designed specifically for Random Forest. We implemented CPI using the ‘permimp’ package, as detailed elsewhere^78^. Briefly, the original permutation importance^25^ shuffled the observations of one feature at a time while holding the target and other features in the same order. Researchers then examined decreases in prediction accuracy in the out-of-bag observations due to the permutation of some features. Stronger decreases are then assumed to reflect the importance of such features. However, this method has shown to be biased when there are correlated features^79^. CPI corrected for this bias by constraining the feature permutation to be within partitions of other features, which was controlled by the threshold ‘s’ value. We used the default s value at .95, which assumed dependencies among features^78^.

SHAP^38^ is a model-agnostic explainer, designed to explain the contribution of each feature to the prediction from any machine learning models via Shapley values^80^. We implemented SHAP using the ‘fastshap’ package (https://bgreenwell.github.io/fastshap/). Based on the cooperative game theory, a Shapley value^80^ quantifies a fair distribution of a payout to each player based on his/her contribution in all possible coalitions where each coalition includes a different subset of players. When applying Shapley values to machine learning, researchers treat each feature as a player in a game, a model output as a pay out and subsets of features as coalitions. Shapley values reflect the weighted differences in a model output when each feature is included vs. not included in all possible subsets of features. SHAP^38^ offers a computationally efficient approach to estimate Shapley values.

### Testing whether the brain-based predictive models mediated the relationships of the behaviourally derived *g*-factor with socio-demographic, psychological and genetic factors

Using leave-one-site-out cross-validation, we built three predictive models for the *g*-factor from (1) multimodal MRI (see above under “Out-of-site Predictive Ability of Multimodal MRI”), (2) key socio-demographic and psychological factors and (3) a polygenic score. This resulted in three types of predicted values of the *g*-factor of unseen children at each held-out data collection site: brain-based *g*-factor, socio-demography-and-psychology-based *g*-factor and gene-based *g*-factor, respectively. We then test if the brain-based *g*-factor mediated the relationship that the behaviourally derived *g*-factor had with the socio-demography-and-psychology-based and gene-based *g*-factors.

### Key socio-demographic and psychological factors

We performed leave-one-site-out cross-validation to build ‘socio-demographic-and-psychological-based’ predictive models. These models predicted the behaviourally derived *g*-factor from key socio-demographic and psychological factors on the baseline data, similar to using leave-one-site-out cross-validation to create the ‘brain-based’ predictive models above. This enabled us to extract predicted values of the *g*-factor based on key socio-demographic and psychological factors at each held-out site, called socio-demography-and-psychology-based *g*-factor. Here we applied a similar modelling approach with leave-one-site-out cross-validation for multimodal MRI, except that we used only one layer of Elastic Net tuned with 200 levels of the penalty (from 10^-10 to 10) and 11 levels of the mixture (from 0 to 1). For pre-processing, we first imputed missing values of the categorical features via mode replacement and then converted them to dummy variables. We next normalised these dummy variables and all numerical features and the behaviourally derived *g*-factor. At the last pre-processing step, we used k-nearest neighbour with five neighbours to impute the missing values of the normalised, numerical features.

Key socio-demographic and psychological factors included 70 features^30^ collected at the baseline (9-10 years old): child’s mental health based on symptom scales in Child Behavioral Checklist^81^ (8 features), primary caretaker’s mental health based on personal strengths and symptom scales in Aseba Adult Self Report^81^ and General Behavior Inventory-Mania^82^ (9 features), child’s personality based on Behavioral Inhibition System/Behavioral Activation System^83^ and the UPPS-P Impulsive Behavior Scale^84^ (9 features), child’s sleep problems based on Sleep Disturbance Scale^85^ (8 features), child’s physical activities based on Youth Risk Behavior Survey^86^ (4 features), child screen use^87^ (4 features), parental use of alcohol, tobacco and marijuana after pregnancy based on Developmental History Questionnaire^88, 89^ (3 features), child developmental adversity (prematurity, birth complications and pregnancy complications) based on Developmental History Questionnaire^88, 89^ (3 features), child socio-demographics^90^ including sex, race, bilingual use^91^, parental marital status, parental education, parental income, household size, economic insecurities, area deprivation index^92^, lead risk^93^, crime reports^94^, neighbourhood safety^95^ and school environment, involvement and disengagement^96^ (17 features) and child social interactions based on Parental Monitoring Scale^97^, Child Report of Behavior Inventory^98^, Strengths and Difficulties Questionnaire^99^ and Moos Family Environment Scale^100^ (5 features).

### Polygenic scores

To extract predicted values of the *g*-factor based on genetics, we used polygenic scores (PGS) for adult cognitive ability^35^. The ABCD study provided details on genotyping elsewhere^101^. Briefly, the study took saliva and whole blood samples and genotyped them using Smokescreen™ Array. The ABCD applied quality control based on calling signals and variant call rates, ran the Ricopili pipeline and imputed the data with TOPMED. We excluded data from problematic plates and with a subject-matching issue, identified by the ABCD. We further quality controlled the data as follows. Firstly, we removed individuals with minimal or excessive heterozygosity. We also excluded SNPs based on minor allele frequency (<5%) and violations of Hardy-Weinberg equilibrium (P < 1E-10). We limited the analysis to “unrelated individuals” as defined by individuals with low genetic relatedness (more than 3rd-degree relative pairs; identical by descent (IBD) ≥0.0422).

We defined alleles associated with the *g*-factor as those related to cognitive ability in a large-scale discovery GWAS sample of European ancestry (N = 257,841)^35^. Given the lower predictive performance of PGS when the ancestry of a sample does not match with that of the discovery GWAS sample,^102^ we restricted all analyses related to PGS to children of European ancestry^102^. We considered children to be genetically similar to the ancestry reference if they were within four standard deviations of the mean of the top four principal components (PCs) of the super-population individuals in the 1000 genomes Phase 3 reference genotypes^103^.

We used the Pthreshold PGS approach where we defined risk alleles as those associated with cognitive ability within the discovery GWAS sample^35^ at 10 different thresholds from p<.5 to .00000001 (referred to as PGS thresholds). The final sample for PGS included 4,814 children (2,263 females; M_age_= 9.94 (SD=.61) years). We computed PGS as the Z-scored, weighted mean number of linkage independent risk alleles. While the *g*-factor was significantly related to the PGS of cognitive ability across thresholds (Figure 7 below), the relationship at the p<.01 PGS threshold was the numerically strongest (r=.21, p<.001 [CI95%=.18-.24]). Accordingly, we focused our analyses using the p<.01 PGS threshold and treated the the PGS at this threshold as our gene-based *g*-factors.

### Mediation analyses

To examine the extent to which brain-based, stacked predictive models of the *g*-factor accounted for the relationship between the behaviourally derived *g*-factor and the socio-demographic, psychological and genetic factors, we applied mediation analyses^104^. In these mediation analyses, we treated (i) the brain-based *g*-factor as the mediator, (ii) the socio-demography-and-psychology-based and gene-based *g*-factors as the independent variables and (iii) the behaviourally derived *g*-factor as the dependent variable. Note the behaviourally derived *g*-factor was computed based on the CFA models in the training data, which were later applied to each held-out site. While the behaviourally derived *g*-factor was a latent variable, it represented the only “observed” value here since the other three *g*-factors (brain-based, socio-demography-and-psychology-based and gene-based) were “predicted” values from predictive models.

We conducted three mediation analyses. The first analysis only used the socio-demography-and-psychology-based *g*-factor as the independent variable. The second analysis only used the gene-based *g*-factor as the independent variable. The third analysis used both the socio-demography-and-psychology-based and gene-based *g*-factors as the independent variables, simultaneously in the same model. To control for population stratification in genetics, we also included four PCs as control variables in the mediation analyses involving the gene-based *g*-factor.

To implement the mediation analyses, we used structural equation modelling (SEM) with 5,000 bootstrapping iterations via lavaan^53^. We specifically calculated the *indirect effects* to show whether the relationships between the behaviourally derived *g*-factor and the socio-demography-and-psychology-based and gene-based *g*-factors were significantly explained by the brain-based *g*-factor. Along with the indirect effects, we also computed the *proportion mediated* to demonstrate the proportion of variance accounted for by the brain-based *g*-factor.

### Data and Code Availability

We used publicly available data provided by the Adolescent Brain Cognitive Development (ABCD) Study (https://abcdstudy.org), held in the NIMH Data Archive (https://nda.nih.gov/abcd/).

We uploaded the R analysis script and detailed outputs for predictive modelling: https://narunpat.github.io/GFactorModelingABCD3/GFactorModelingABCD3.html and mediation analyses: https://narunpat.github.io/GFactorModelingABCD3/MediationSocDemPsycPGSBrainABCD3. html .

## Results

### How Robust Are the Factor Scores of the *g*-factor Based on the 2^nd^-order Model?

Based on our confirmatory factor analysis (CFA), the 2^nd^-order model of the *g*-factor showed a good fit: (a) scaled, robust Comparative Fit Index (CFI) =.995, (b) scaled, robust Tucker-Lewis Index (TLI) =.988, (c) scaled, robust Root Mean Square Error of Approximation (RMSEA) = .029 (90%CI=.015-.043) and (d) robust Standardized Root Mean Square Residual (SRMR) = .014. The *g*-factor latent variable of the 2^nd^-order model also had high internal consistency: OmegaL2=.78.

See Supplementary Appendix 1 and 2 for a more detailed CFA of the *g*-factor. In brief, firstly, the 2^nd^-order model had better fit indices than the single-factor model. Additionally, factor scores of the *g*-factor from the 2^nd^-order model, the single-factor model, and the mixture between EFA and CFA models were similar to each other at high magnitude (Pearson’s *rs* ≥ .987). Accordingly, the choice of *g*-factor models had only minimal effects on the estimation of the factor scores for the *g*-factor, and thus our brain-based predictive models should be generalisable to the factor scores of different *g*-factor CFA models beyond the 2^nd^-order model. Lastly, the factor scores estimated from the 1^st^-layer training data were similar to the factor scores estimated from the full baseline data at high magnitude (Pearson’s *rs* > .997), indicating the stability of the factor scores used.

### How Robust Are the Brain-Based Predictive Models?

#### 1) Out-of-sample Predictive Ability of Multimodal MRI

For hyperparameter-tuning results, see Supplementary Appendix 3. Table 1 and Figure 2 summarise the out-of-sample predictive ability of multimodal MRI for both baseline and follow-up samples. Performance of Stacked All, Stacked Complete and Stacked Best was among the top with Pearson’s r over .4 and R^2^ around .19. Importantly, the superior performance of stacked models was found across baseline and follow-up test sets at a similar magnitude, suggesting their longitudinal stability. Note that given that the N-Back task-based fMRI had the highest performance among modality-specific models, we set the missing values of the Stacked Best to be the same as those of the N-Back task-based fMRI. Moreover, the opportunistic stacking^24^ algorithm that led to the stacked model with at least one modality present, Stacked All, was robust against missing values as the performance of Stacked All was similar to that of the stacked model with all modalities present, Stacked Complete.

**Figure 2.**
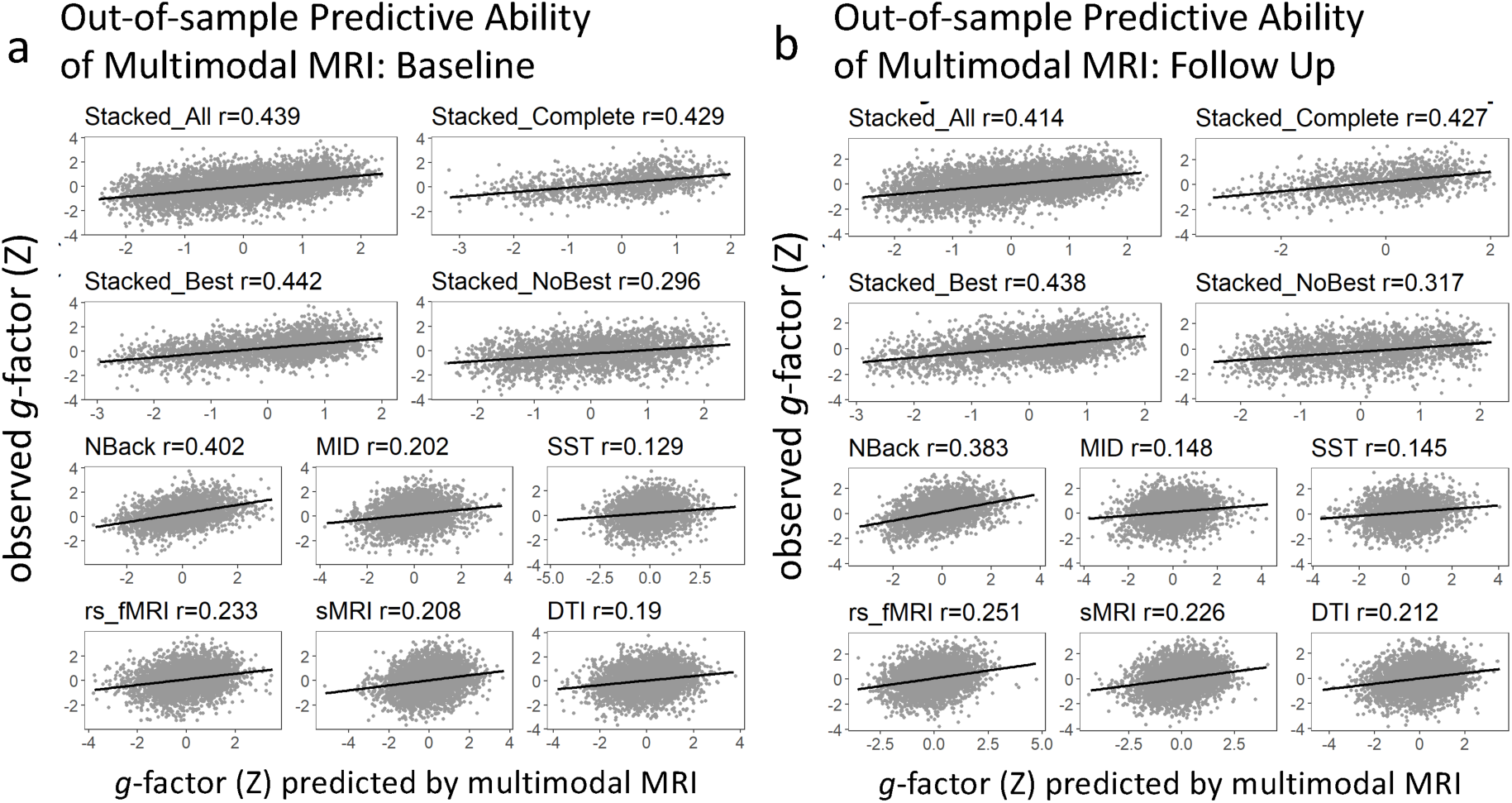
Out-of-sample Predictive Ability of Multimodal MRI as a Function of Modalities in the Test Sets for Baseline (2A) and Follow-Up (2B) Samples. Stacked All required the test data with at least one modality present. Stacked Complete required the test data with all modalities present. Stacked Best had the same missing values with the modality with the best prediction (N-Back task-based fMRI). Stacked No Best did not have any test data from the modality with the best prediction and had at least one modality present.

**Table 1.** Out-of-sample and out-of-site predictive ability of multimodal MRI. r = Pearson’s correlation; R^2^ = coefficient of determination; MAE = mean absolute error; RMSE = root mean squared error.

| Models | $r$ | $R^2$ | MAE | RMSE |
| --- | --- | --- | --- | --- |
| Stacked_All | 0.439 | 0.191 | 0.699 | 0.895 |
| Stacked_Complete | 0.429 | 0.183 | 0.61 | 0.78 |
| Stacked_Best | 0.442 | 0.195 | 0.62 | 0.798 |
| Stacked_NoBest | 0.296 | 0.085 | 0.783 | 0.987 |
| NBack | 0.402 | 0.072 | 0.664 | 0.857 |
| SST | 0.129 | -0.033 | 0.744 | 0.95 |
| MID | 0.202 | 0.013 | 0.738 | 0.944 |
| rs_fMRI | 0.233 | 0.042 | 0.749 | 0.955 |
| sMRI | 0.208 | 0.04 | 0.763 | 0.969 |
| DTI | 0.19 | 0.033 | 0.757 | 0.972 |

Out-of-sample predictive ability of multimodal MRI: follow-up samples
| Models | $r$ | $R^2$ | MAE | RMSE |
| --- | --- | --- | --- | --- |
| Stacked_All | 0.414 | 0.166 | 0.719 | 0.913 |
| Stacked_Complete | 0.427 | 0.168 | 0.651 | 0.829 |
| Stacked_Best | 0.438 | 0.175 | 0.666 | 0.846 |
| Stacked_NoBest | 0.317 | 0.1 | 0.794 | 1 |
| NBack | 0.383 | 0.118 | 0.687 | 0.875 |
| SST | 0.145 | -0.004 | 0.76 | 0.961 |
| MID | 0.148 | -0.003 | 0.757 | 0.955 |
| rs_fMRI | 0.251 | 0.055 | 0.754 | 0.954 |
| sMRI | 0.226 | 0.049 | 0.764 | 0.965 |
| DTI | 0.212 | 0.045 | 0.771 | 0.978 |

Mean (SD) of out-of-site predictive ability of multimodal MRI
| Models | $r$ | $R^2$ | MAE | RMSE |
| --- | --- | --- | --- | --- |
| Stacked | 0.46 (0.057) | 0.21 (0.052) | 0.698 (0.023) | 0.888 (0.029) |
| NBack | 0.408 (0.069) | 0.167 (0.055) | 0.718 (0.028) | 0.91 (0.031) |
| MID | 0.227 (0.096) | 0.05 (0.05) | 0.772 (0.021) | 0.973 (0.025) |
| SST | 0.139 (0.071) | 0.019 (0.024) | 0.783 (0.014) | 0.988 (0.012) |
| rs_fMRI | 0.255 (0.061) | 0.064 (0.03) | 0.765 (0.015) | 0.966 (0.016) |
| sMRI | 0.248 (0.092) | 0.061 (0.046) | 0.763 (0.024) | 0.967 (0.024) |
| DTI | 0.223 (0.076) | 0.049 (0.037) | 0.766 (0.016) | 0.974 (0.019) |

Figure 3 shows the proportion of missing data in the two test sets. sMRI had the lowest missing observations, while the three task-based fMRI data had the highest. Missing observations in Stacked All were around 3-6%, while those in Stacked Complete were up to 78.79%. Figure 3 also shows the differences in the *g*-factor between participants with vs. without missing values for each model in the two test sets. Participants with missing values had a significantly lower g-factor than those without missing values, as indicated by Welch’s t-test, for all models, except for Stacked No Best, which showed the opposite direction. Yet, numerically these differences in the g-factor were weaker in magnitude in the Stacked All than in other models with high predictive performance (such as the N-Back task-based fMRI and Stacked Complete) as indicated by Cohen’s d. Accordingly, by imputing the data via the opportunistic stacking^24^, we were able to include more participants, and thus, less likely to exclude participants with a lower *g*-factor.

**Figure 3.**
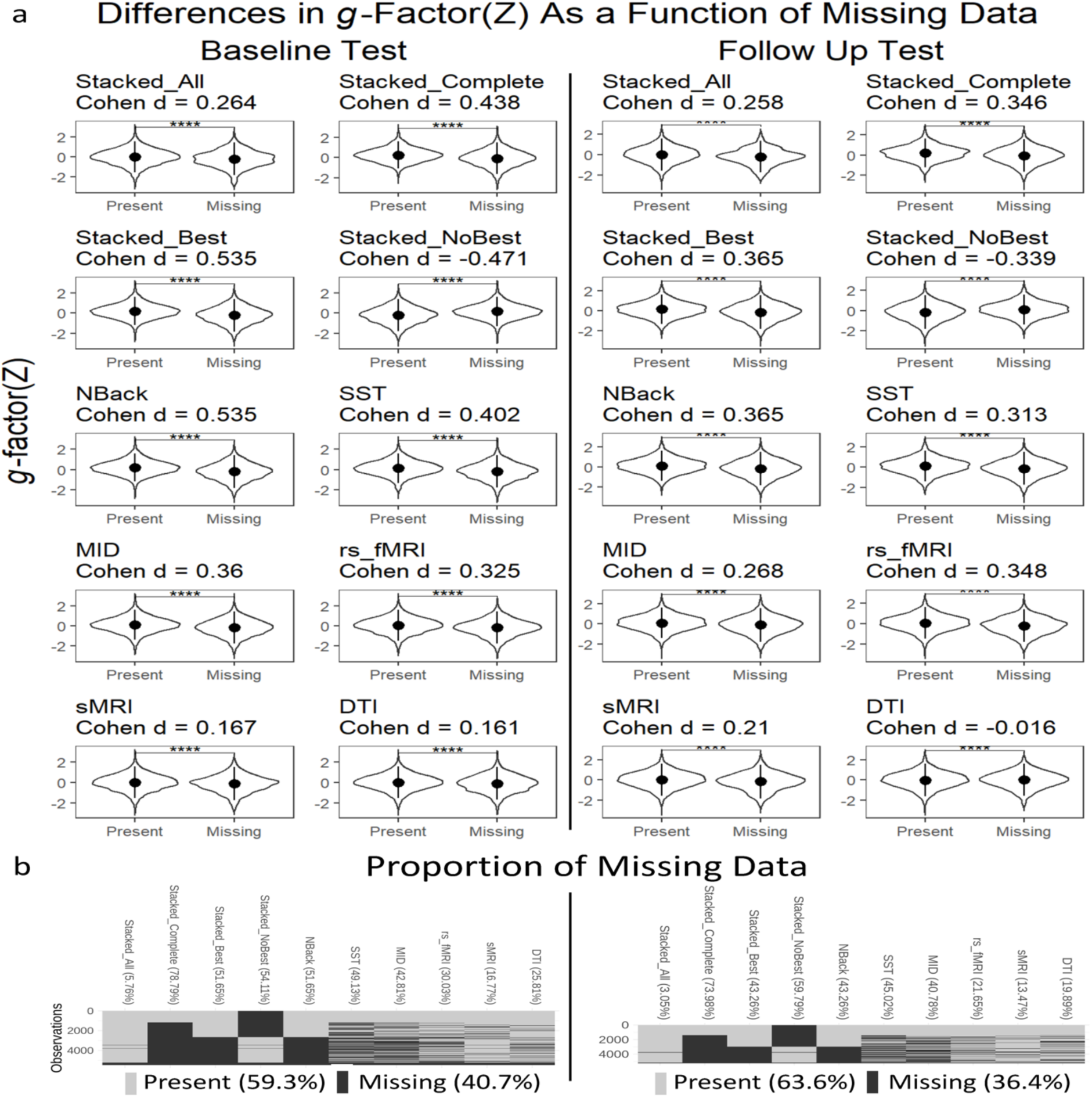
Missing Values in Each Predictive Model in the Baseline and Follow-Up Test Sets. 3A shows the differences in the g-factor between participants with vs. without missing values for each predictive model in the two test sets. **** indicates p-value < .001 based on Welsh’s t-test. Positive Cohen’s d indicates that participants without missing values had a higher g-factor than participants without missing values. Dot and line are the mean and standard deviation x 2 of the g-factor, respectively. 3B shows the proportion of missing data for each predictive model in the two test sets.

#### 2) Comparing Out-of-sample Predictive Ability of Multimodal MRI between the Stacked Model and N-Back Task-Based fMRI

N-Back task-based fMRI provided the best out-of-sample predictive ability for both baseline and follow-up test sets, relative to other modality-specific models. Figure 4 compared the predictive ability between the Stacked Best and N-Back task-based fMRI using bootstrapped differences. The Stacked Best had significantly higher performance in both baseline and follow-up test sets, reflected by higher Pearson’s *r* and *R*^2^ and lower MAE and RMSE. This indicates the boost in predictive performance when multiple modalities were integrated, at around 12% for the baseline data and 6% for the follow-up data. Accordingly, the stacked model performed better than the best single modality.

**Figure 4.**
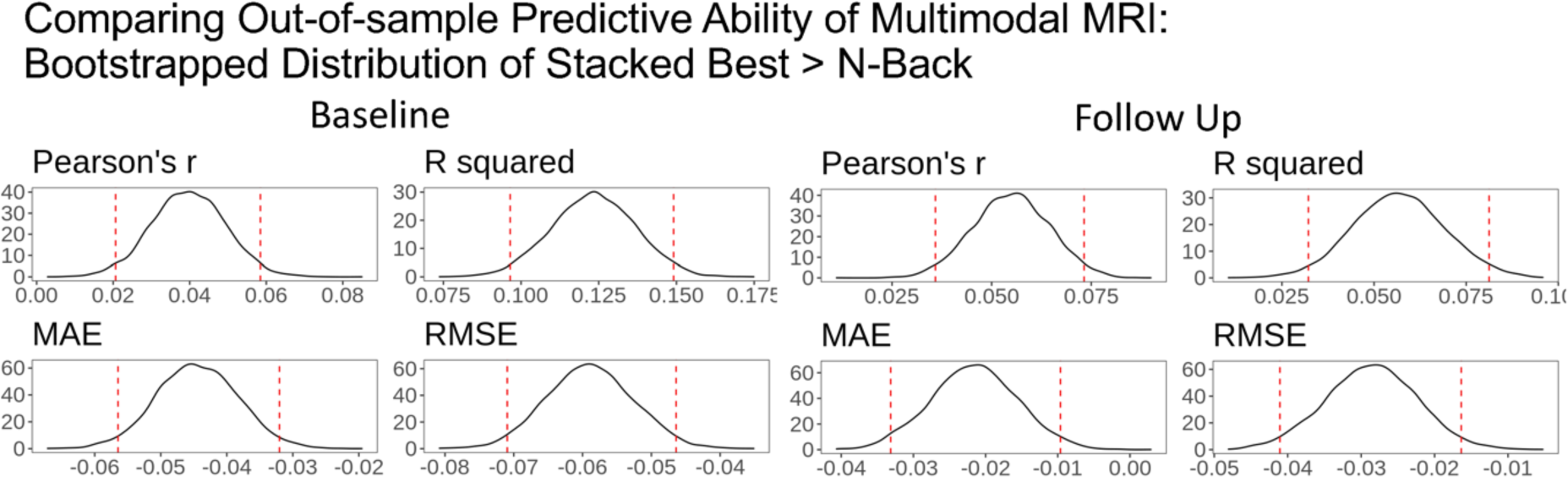
Comparing Out-of-sample Predictive Ability of Multimodal MRI between Stacked Best and the Model with the Best Modality (N-Back Task-Based fMRI). Here we separately applied bootstrapping on the baseline and follow-up test sets. At each of 5,000 iterations, we computed performance indices (including r, R^2^, MAE, and RMSE) of Stacked Best and N-Back Task-Based fMRI models and subtracted performance indices of N-Back Task-Based fMRI from that of Stacked Best. Dotted lines indicate 95% confidence intervals. R squared = coefficient of determination; MAE = mean absolute error; RMSE = root mean squared error.

#### 3) Out-of-site Predictive Ability of Multimodal MRI

Based on leave-one-site-out cross-validation, the out-of-site predictive ability of the stacked model was highest, explaining on-average 21% (SD =5.2) of the variance in the *g*-factor across 21 sites (see Table 1 and Figure 5). This confirmed the generalisability of the stacked model and ensured its use for subsequent mediation analyses.

**Figure 5.**
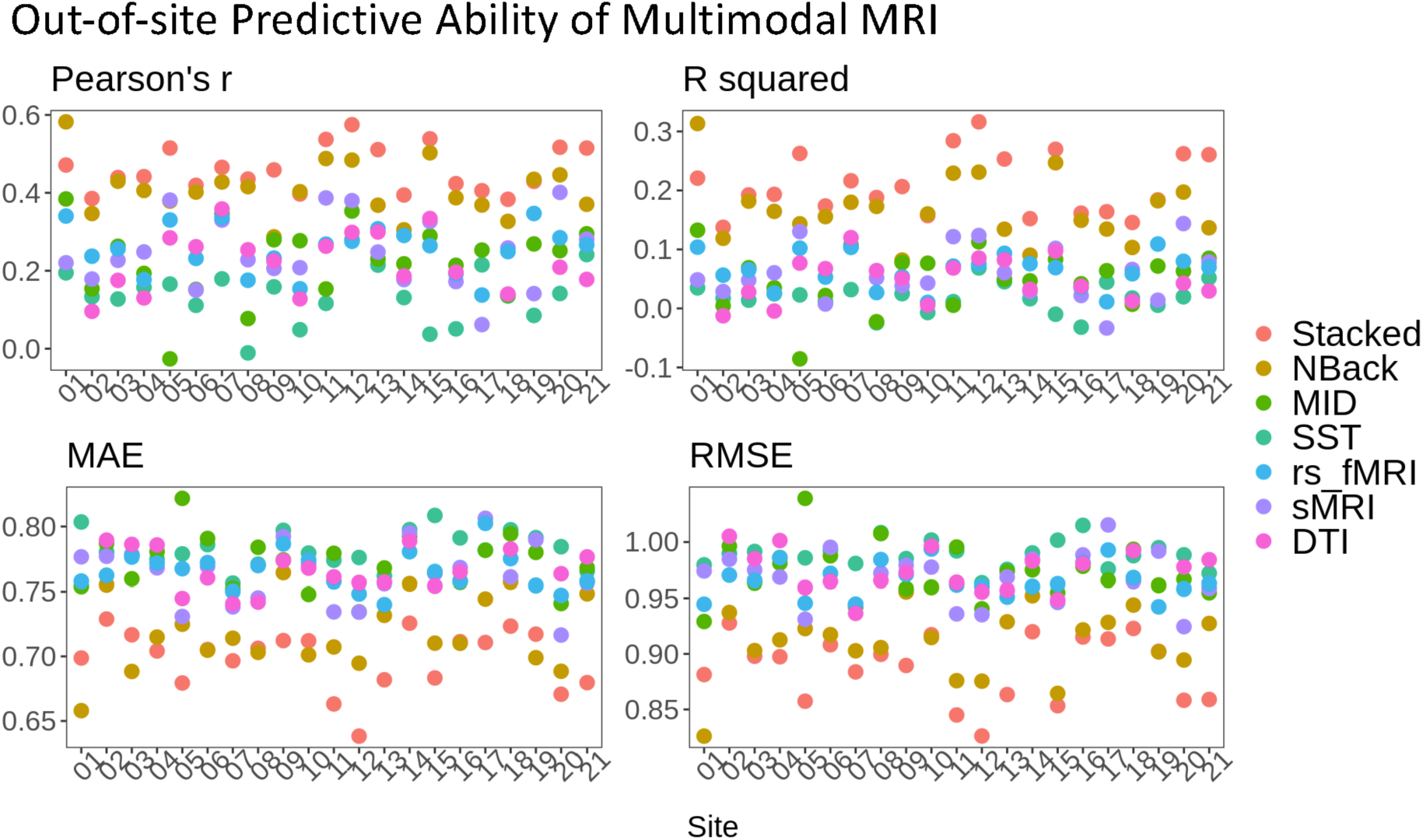
Out-of-site predictive ability of multimodal MRI via leave-one-site-out cross-validation. We evaluated out-of-site predictive ability between predicted vs. observed g-factor in the held-out site. Note that DTI data were not available from 3 sites (sites 1, 17, and 19). R squared = coefficient of determination; MAE = mean absolute error; RMSE = root mean squared error.

### Feature Importance of Multimodal MRI Models

Figure 6 shows the feature importance of both the modality-specific and stacked models. For the modality-specific models, we applied eNetXplorer^36^ to show brain features that significantly (empirical p <.05) contributed to the prediction. For N-Back task-based fMRI, the *g*-factor prediction was driven by activity in areas, such as the precuneus, sulcus intermedius primus, superior frontal sulci, and dorsal cingulate. For MID task-based fMRI, the prediction was driven by activity in several areas in the parietal, frontal and temporal regions. For SST, the prediction was contributed by activity in areas such as the supramarginal gyrus and inferior precentral sulcus. For rs-fMRI, the prediction was driven by connectivity within cinguloparietal and sensory-motor-hand as well as between networks that were connected with frontoparietal, default-mode, and sensory-motor-hand networks. For sMRI, the prediction was driven by the volume/thickness at several areas, such as the insula, middle frontal gyrus, and lingual sulcus. For DTI, the prediction was driven by FA at several white matter tracts, such as the superior longitudinal fasciculus, forceps minor, uncinate and parahippocampal cingulum. For the stacked model, we applied the Conditional Permutation Importance (CPI)^37^ and SHapley Additive exPlanations (SHAP)^38^ to examine which of the modalities contributed strongly to the prediction. CPI and SHAP provided similar results. N-Back task-based fMRI by far had the highest importance score.

**Figure 6.**
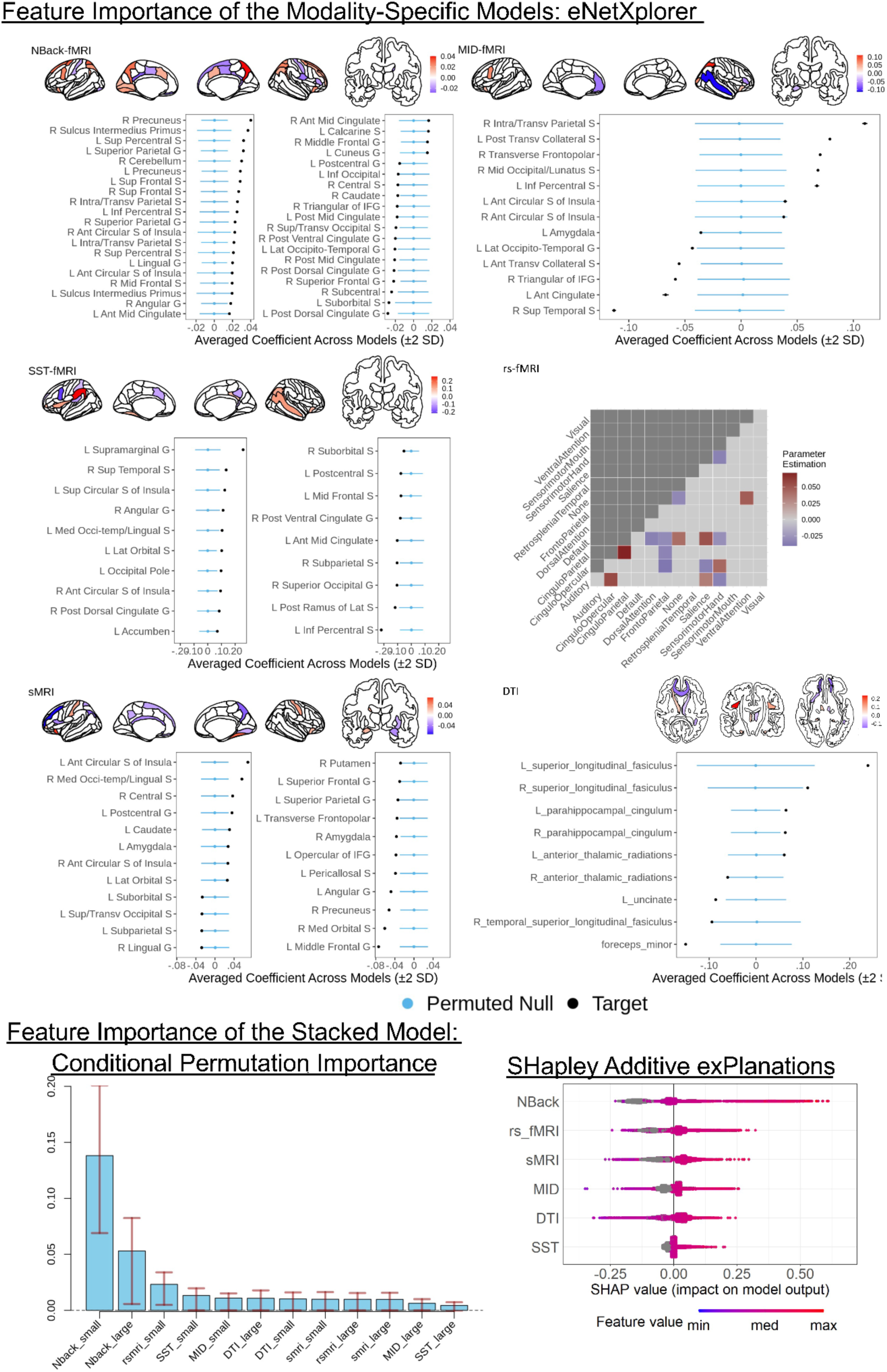
Feature importance of the modality-specific and stacked models. For the modality-specific models, we applied eNetXplorer^36^ permutation and only plotted brain features with empirical p < .05. For the stacked model, we applied Conditional Permutation Importance (CPI)^78^ and SHapley Additive exPlanations (SHAP)^38^. Both CPI and SHAP were computed based on the second-layer training set. Error bars in the CPI plot show an interval between .25 and .75 quantiles of the CPI for each tree in the Random Forests. The “_large” and “_small” suffixes indicate whether the missing values were coded as a large (1000) or small (−1000) number, respectively. For SHAP, we combined Shapley values across the two coded features of the same modality. We then ranked the modalities according to the absolute value of SHAP; the highest one was N-Back task-based fMRI. Note the grey colour indicates observations with a missing value (coded as 1000 or −1000). Sup=Superior; Ant=Anterior; Lat=lateral; Med=Medial; S=Sulcus; G=Gyrus; IFG=Inferior Frontal Gyrus; L=left; R=right.

### Did the brain-based predictive models mediate the relationships of the behaviorally derived *g*-factor with socio-demographic, psychological and genetic factors?

#### Key socio-demographic and psychological factors

Based on leave-one-site-out cross-validation, socio-demographic and psychological factors explained on-average 29.7% (SD =8.1) of the variance in the behaviourally derived *g*-factor across sites (see Figure 7). The top features in the Elastic-Net models that had the magnitude of their standardised coefficients over .1 included parents’ education and income along with child’s attention and social problems as well as extracurricular activities.

**Figure 7.**
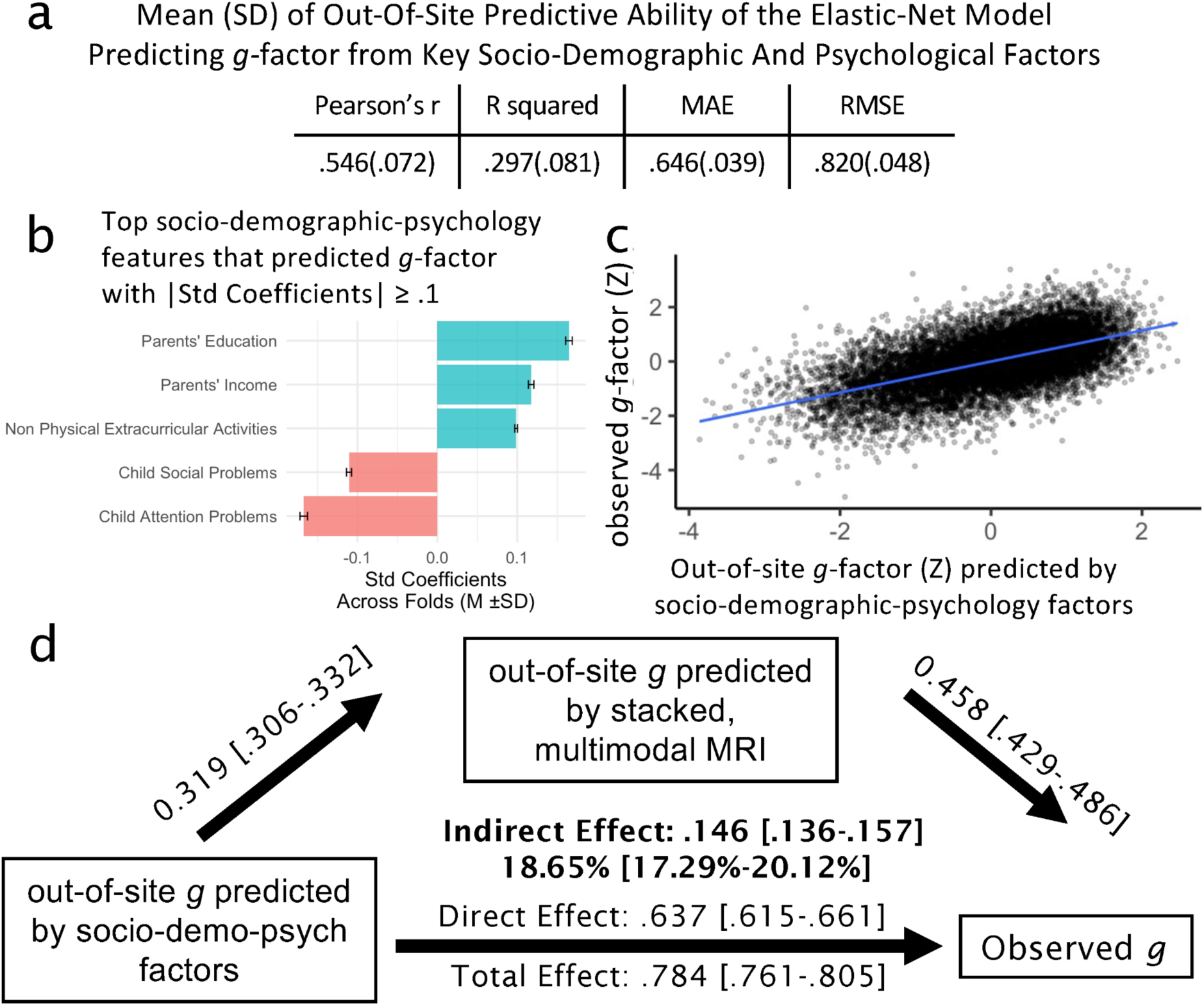
Key socio-demographic and psychological factors. 7a shows the out-of-site predictive ability of the Elastic-Net model predicting the g-factor from key socio-demographic and psychological factors, based on leave-one-site-out cross-validation. 7b shows the top socio-demographic and psychological features with the magnitude of standardised coefficient over .1 based on the Elastic-Net model. Blue indicates a positive relationship while red indicates a negative relationship. 7c shows a scatter plot between out-of-site predicted values of the g-factor based on key socio-demographic and psychological factors and the observed (i.e., real) values of the g-factor. 7d shows a mediation analysis where (1) the socio-demography-and-psychology-based g-factor (the out-of-site predicted values of the g-factor based on the key socio-demographic and psychological factors at all held-out sites) is the independent variable, (2) the brain-based g-factor (the out-of-site predicted values of the g-factor of the stacked model based on multimodal MRI data at all held-out sites) is the mediator and (3) the behaviourally derived g-factor (the observed g-factor) is the dependent variable. % under the indirect effect indicates proportion mediated. [ ] indicates a 95% confidence interval based on bootstrapping. R squared = coefficient of determination; MAE = mean absolute error; RMSE = root mean squared error.

#### Polygenic scores (PGS)

Figures 8a,b show the relationship between the behaviourally derived *g*-factor and the PGS of cognitive ability at different thresholds. While the behaviourally derived *g*-factor was significantly related to the PGS of cognitive ability across thresholds, the relationship at the *p*<.01 PGS threshold was the numerically strongest (*r*=.21, p<.001 [CI95%=.18-.24]). Accordingly, we used PGS at the *p*<.01 PGS threshold as the gene-based *g*-factor for the mediation analyses.

**Figure 8.**
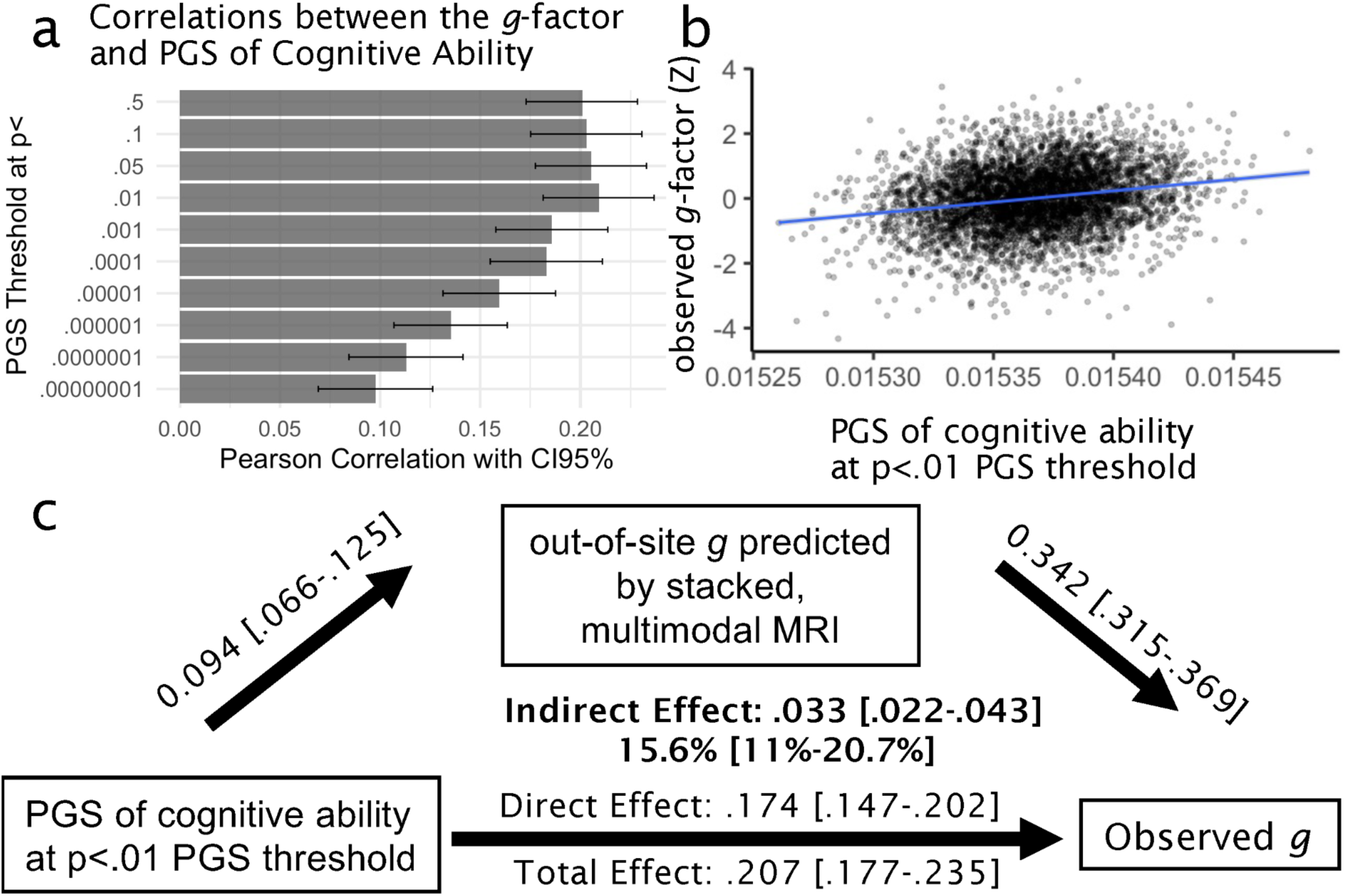
Polygenic Scores (PGS) of cognitive abilities. 8a shows Pearson’s correlations between the g-factor and PGS of cognitive abilities at different PGS thresholds. 8b shows a scatter plot between the PGS of cognitive abilities at the p<.01 PGS threshold and the observed (i.e., real) values of the g-factor. 8c shows a mediation analysis where (1) gene-based g-factor (the PGS of cognitive ability at the p<.01 PGS threshold) is the independent variable, (2) the brain-based g-factor (the predicted values of the g-factor of the stacked model based on multimodal MRI data at all held-out sites) is the mediator and (3) the behaviourally derived g-factor (the observed g-factor) is the dependent variable. Not shown in the figure are four principal components included as the control variables to adjust for population stratification. % under the indirect effect indicates proportion mediated. [ ] indicates a 95% confidence interval based on bootstrapping.

#### Mediation analyses

We tested whether brain-based *g*-factor mediated the relationships between the behaviourally derived *g*-factor and socio-demography-and-psychology-based and gene-based *g*-factors. We found significant indirect effects (1) when the socio-demography-and-psychology-based *g*-factor was the sole independent variable (Figure 7d proportion mediated = 19.1%), (2) when the gene-based *g*-factor was the sole independent variable (Figure 8c, proportion mediated = 15.6%) and (3) when both socio-demography-and-psychology-based *g*-factor (Figure 9, proportion mediated = 15%) and gene-based *g*-factor (Figure 9, proportion mediated = 10.75%) were the covaried independent variables.

**Figure 9.**
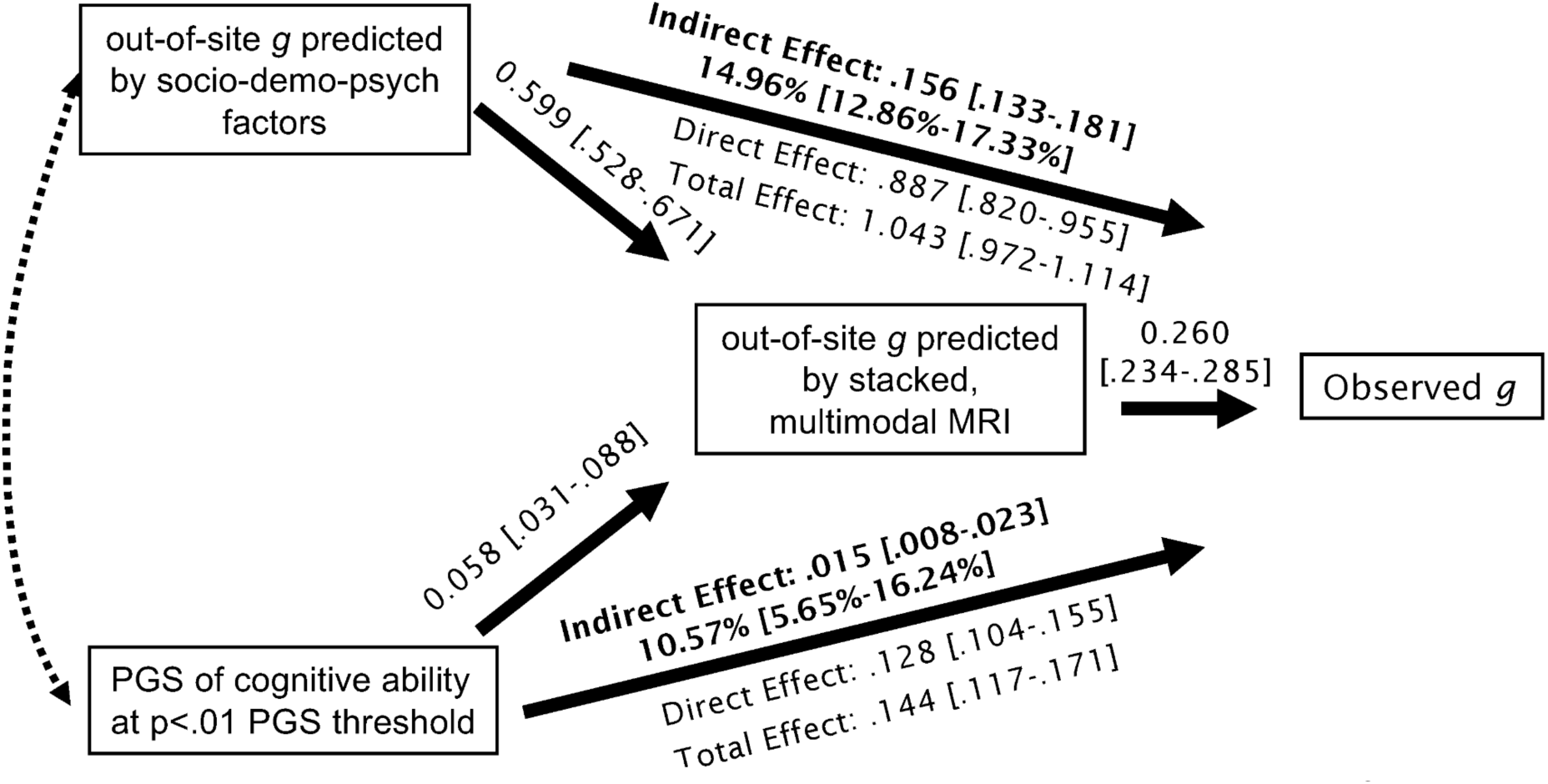
Mediation analysis with both key socio-demographic and psychological factors as well as genetic factors as independent variables. Specifically, this model treated (1) the socio-demography-and-psychology-based g-factor (the out-of-site predicted values of the g-factor based on the key socio-demographic and psychological factors at all held-out sites) and (2) the gene-based g-factor (the PGS of cognitive ability at the p<.01 PGS threshold) as two separate independent variables. It treated the brain-based g-factor (the predicted values of the out-of-site predicted values of the g-factor of the stacked model based on multimodal MRI data at all held-out sites) as the mediator and the behaviourally derived g-factor (the observed g-factor) as the dependent variable. Not shown in the figure are four principal components included as the control variables to adjust for population stratification. % under the indirect effect indicates proportion mediated. [ ] indicates a 95% confidence interval based on bootstrapping. The dotted, double arrowed line indicates covariation between the two independent variables.

## Discussion

Following the Research Domain Criteria’s (RDoC) integrative approach for cognitive abilities^1^, we aimed to develop brain-based predictive models that can a) improve our current ability to predict children’s cognitive abilities and b) account for the relationships between cognitive abilities and socio-demographic, psychological and genetic factors. Here we showed that incorporating data from different MRI modalities into stacked models substantially improved our ability to predict cognitive abilities, operationalised as the behaviourally derived *g*-factor. Our brain-based, stacked predictive models were stable across years and generalisable to different sites while being able to handle missing values. Moreover, we showed that the brain-based, stacked models significantly, albeit partially, mediated the relationships of the behaviourally derived *g*-factor with socio-demographic, psychological and genetic factors. Thus, our brain-based predictive for children’s *g*-factor demonstrated construct validity according to the RDoC framework^1, 6, 7, 31^.

### The brain-based, stacked predictive models for the *g*-factor were 1) predictive, 2) longitudinal stable, 3) robust against missing values and 4) explainable

We developed longitudinal predictive models for children’s *g*-factor from MRI data of different modalities. We built models from the baseline MRI data and tested them on unseen children at the same age and two years older. We found similar predictive ability across these two test sets for all modality-specific and stacked models. That is, the models that had high out-of-sample prediction on same-age children also had high out-of-sample prediction on older children, suggesting the longitudinal stability of MRI for many modalities. The best model across all performance indices (Pearson’s *r*, *R*^2^, MAE, and RMSE) was the stacked model that incorporated all six modalities, which was followed closely by the N-back task-related fMRI model. Apart from the SST task-related fMRI model, other models (including the MID tasked-related, rs-fMRI, sMRI, and DTI) performed moderately well. We also found a similar magnitude for out-of-site predictive ability based on leave-one-site-out cross-validation, suggesting the generalisability of MRI not only across ages but also across data collection sites. Overall, the stacked model partially predicted the children’s *g*-factor at around 20% of the variance. This made the stacked model the most generalisable to out-of-sample, out-of-site children and the most longitudinally stable.

Beyond generalisability across ages and sites, the stacked model based on opportunistic stacking^24^ also allowed us to handle missingness in the data. This is especially important for children’s MRI data given high levels of noise in certain modalities^22^. If we were to use data only from children with all modalities present (i.e., the Stacked Complete), the model would not apply to around 80% of the children. The opportunistic stacking allowed us to use the data as long as one modality was present (i.e., the Stacked All), leaving the exclusion to just around 5%. Importantly, the predictive performance of Stacked Complete and Stacked All were both relatively high, ensuring the ability of opportunistic stacking to deal with the missing data. Furthermore, handling missingness in the data via opportunistic stacking also heightened the chance of including participants with a wider range of the *g*-factor, including those with a lower *g*-factor who usually had missingness in the MRI data (perhaps due to high movement artifacts^22^). Moreover, in the case when the best modality was not available, using the stacked model (i.e., the Stacked No Best) could be helpful. While the predictive ability of the Stacked No Best was not as strong as the Stacked Complete, Stacked All and Stacked Best, its performance measures of variance (Pearson’s r and R squared) appeared stronger in magnitude than any other non-optimal modalities by themselves. Accordingly, in settings where not all of the modalities are available, researchers/practitioners can still take advantage of the boosted predictive ability of the stacked models over unimodal models.

The stacked model improved predictive ability over and above the best modality, which was the N-Back task-based fMRI. This is based on bootstrapping distributions of the differences in performance indices between the N-Back task-based fMRI and the stacked model with the same participants (i.e., the Stacked Best). Our finding is consistent with previous studies showing the enhanced predictive power of the stacked model^19, 24^. Yet, it is important to note that, while the improvement in performance was statistically significant, the magnitude of this improvement was somewhat modest. For instance, in the case of the baseline samples, the Stacked Best led to *r* = .442 and *R*^2^ =.195, which was improved from the N-Back task-based fMRI at *r* = .402 and *R*^2^ =.072, rendering the improvement at around r ∼.04 and *R*^2^ ∼.123. Accordingly for researchers who have access to all MRI modalities and several fMRI tasks, including the N-Back task, using the stacked model should provide the best possible performance for predicting the *g*-factor. However, if resources are constrained, the next best option would be using the N-Back task-based fMRI along with other modalities that are available.

In addition to predictability, our machine learning framework allowed for easy-to-explain models, highlighting the neurobiological bases of children’s *g*-factor. Explainability is used in a specific machine-learning sense^105^, referring to the extent to which a technique applied allows us to explain the contribution of each brain feature to the prediction. Here, conditional permutation importance (CPI)^78^ and SHapley Additive exPlanations (SHAP)^38^ allowed us to infer that prediction from the stacked model was driven primarily by N-back task-related fMRI. This indicates the important role of working memory. eNetXplorer permutation^36^ further showed us that contribution from fMRI activity in the parietal and frontal areas during the N-back task drove the prediction. These areas were similar to the areas previously found in a recent study in adults^106^. Similarly, we also found brain indices from other modalities, from activity during other tasks to the cortical thickness and white matter density, that contributed to the prediction of the *g*-factor, albeit with lower predictive performance.

Unlike previous unimodal^11–18^ and multimodal studies^19, 20^, we were able to compare the ability of task-based fMRI with other modalities in predicting the *g*-factor. We found that one of the three task-based fMRI models, the N-Back, performed exceptionally well. Based on the CPI^78^ and SHAP^38^, the N-Back task-related fMRI appeared to drive the prediction of the stacked model. This finding is consistent with a recent study using adults’ data from the Human Connectome Project, showing superior performance of the N-Back task in predicting the *g*-factor, compared to resting-state fMRI^106^ and other tasks. Showing that task-based fMRI from a certain task could capture cognitive ability across a two-year gap provided a promising outlook for the use of task-based fMRI as a predictive tool. Our finding is contradictory to a more common practice in cognitive neuroscience that usually relies on sMRI^107–109^ or resting-state fMRI^15, 19, 106^ when predicting cognitive abilities. These sMRI and resting-state fMRI studies often result in poorer predictive performance (at *r* < .4) than what was found here. Accordingly, we are in agreement with a recent movement^110^ for studies on individual differences to move from resting-state fMRI and embrace other MRI modalities, including task-based fMRI.

It is important to note that not all fMRI tasks were suitable for predicting certain targets. The N-Back task and SST, for instance, were designed to capture working memory^55, 58^ and inhibitory control^55, 60^, respectively. Accordingly, both should be related to the *g*-factor, especially on memory recall and mental flexibility portions of the *g*-factor. Yet, only the N-Back task showed good predictive ability. This may be due to different cognitive processes in each task (i.e., working memory vs. inhibitory control) or to different task configurations. It is entirely possible, for instance, that the block design used in the N-Back, as opposed to the event-related design used in the SST, allowed the N-Back to have higher predictive power. Accordingly, while task-based fMRI can have high predictive power, systematic comparisons are required in future research to better understand the characteristics of some tasks that make them more suitable for predicting the *g*-factor and other individual differences.

### The brain-based, stacked predictive models for the *g*-factor demonstrated construct validity, according to the RDoC framework^6^

Here we tested the construct validity of the brain-based, stacked predictive models for the *g*-factor according to the RDoC framework^6^. The RDoC proposes that cognitive abilities are affected by socio-demographic and psychological factors^1, 26^. The RDoC also proposes that cognitive abilities as measured by brain differences belong to the same domain as cognitive abilities as measured by gene differences^1, 6, 7, 31^. Accordingly, to satisfy these presuppositions, our brain-based, stacked predictive models for the *g*-factor should be able to capture the relationship between the behaviourally derived *g*-factor and socio-demographic, psychological and genetic factors.

We first built a predictive model of the *g*-factor using 70 socio-demographic and psychological features^30^, resulting in the socio-demography-and-psychology-based *g*-factor. This model had relatively high performance, accounting for around 30% of the *g*-factor. Moreover, the top contributing features are consistent with previous studies, including socio-demographics^27^ (e.g., parents’ education and income) along with children’s mental health^28, 29^ (e.g., attention and social problems) and children’s extracurricular activities^30^. More importantly, our mediation analysis showed that the brain-based *g*-factor captured approximately 19% of the relationship between the behaviourally derived *g*-factor and the socio-demography-and-psychology-based *g*-factor.

As for the genetic factor, we first showed that the polygenic score (PGS) based on adults’ cognitive abilities^35^ was related to children’s *g*-factor, consistent with a recent study^111^. This enabled us to use the PGS of cognitive abilities as the gene-based *g*-factor. Similar to the socio-demography-and-psychology-based *g*-factor, our mediation analysis showed that the brain-based *g*-factor accounted for approximately 16% of the relationship between the behaviourally derived *g*-factor and the gene-based *g*-factor. In fact, mediation from the brain-based *g*-factor was still significant when having both socio-demography-and-psychology-based and gene-based *g*-factors together as independent variables in the model. Altogether, our brain-based, stacked predictive models for the *<u>g</u>*-factor demonstrated the construct validity of cognitive abilities that is in line with the RDoC framework^6^.

### Applications, Limitations, and Disclaimers

For applications, our brain-based predictive models for the *g*-factor facilitate the development of a robust, transdiagnostic research tool for cognition at the neural level in keeping with the RDoC^1^. Cognitive abilities are one of RDoC’s six major transdiagnostic domains^1^, relating to a number of psychiatric disorders^2–4^. Based on RDoC^1^, to improve our understanding of cognitive abilities, we need research tools that allow us to integrate different units of analysis, from behavioural down to neural and genetic levels, and that reflect the influences of socio-demographical and psychological factors across the lifespan^6, 7^. Our brain-based predictive models satisfied many presuppositions of RDoC^1^. Our brain-based predictive models for the *g*-factor were not only longitudinal stable^6, 7^, but they also captured the influences of socio-demographical, psychological and genetic factors on cognitive abilities^1, 6, 7, 31^. In fact, the predictive ability of our brain-based predictive models in capturing the behavioural performance of cognitive tasks was considerably higher than that of PGS (multimodal MRI’s r ∼ .4 and R^2^ ∼ .2 vs. PGS’s r ∼ .21 in our study and R^2^ < .1 in another study^111^), suggesting the potential use of brain-based predictive models for a robust, transdiagnostic, brain-based marker for cognitive abilities.

With opportunistic stacking, those who wish to adapt our brain-based predictive models to compute a transdiagnostic brain-based marker for cognition in their own data, but do not have as many modalities as the ABCD, can still use our models. That is, they can still use the model built from the ABCD and impute missing values of certain modalities to fits with their study. Accordingly, our use of opportunistic stacking provides a scalable and flexible approach for future researchers following the RDoC framework^1^.

Our study is not without limitations. We relied on the ABCD study’s curated, preprocessed data^10, 23, 55^. This provided certain advantages. For instance, given that the curated data provided by the ABCD have already been preprocessed, other studies that wish to apply our model of the *g*-factor to the ABCD data can readily do so without concerns about differences in preprocessing steps. Preprocessed data also enabled us to apply the manual quality control done by the study, a process that required time and well-trained labour^10, 23, 55^.

Preprocessing large-scale multi-modal data ourselves would not only demand significant computer power and time but is prone to error. However, using the preprocessed data only allowed us to follow the choices of processing done by the study. For example, ABCD Release 3 only provided Freesurfer’s parcellation^65, 66^ for task-based fMRI. While this popular method allowed us to explain task-based activity on subject-specific anatomical landmarks, the regions are relatively large compared to other parcellations. Future studies will need to examine if smaller and/or different parcellations would improve predictive performance. Next, our predictive modelling framework was designed to predict the out-of-sample *g*-factor, but not the developmental changes in the *g*-factor, from multimodal MRI. More specifically, we standardised MRI and cognitive data within each age group to satisfy the assumption of our machine-learning algorithms ^72^ and to force behavioural performance from different cognitive tasks onto the same scale. This unfortunately made our predictive models inappropriate for predicting the developmental changes in cognition over years^112^. Future research that aims to capture the developmental changes in cognition would need to employ a different strategy for standardisation^112^.

In terms of important disclaimers, research reporting on cognitive abilities can be misunderstood or misquoted for alien purposes^113^. It is therefore important to clarify the following. First, the fact that measurements taken from the brain were related to cognitive abilities should not be equated with assertions that variability in cognitive abilities is “purely biological”. Here we showed that the predictive model for the *g*-factor based on socio-demographic and psychological variables that were available in the ABCD^30^ already accounted for a larger variance of the *g*-factor (∼30%) than the predictive models based on the brain (∼20%) or genes (<10%^111^). Moreover, our mediation analysis showed that the brain-based predictive models could only account for approximately 19% of the relationship between cognitive abilities and socio-demographic and psychological factors. Accordingly, it is very plausible that social-demographic and psychological circumstances, broadly construed, have at least partial aetiological primacy. Second, it should be clear that social-demographic, psychological and genetic circumstances may not be independent of one another, as suggested by studies on the complex interplay of genes and environments on cognitive abilities over the course of cognitive development^114, 115^. This is shown in our mediation analyses. Here the brain-based *g*-factor showed less proportion mediated for the influences of social-demographic, psychological factors and genes when they were included together in the model, compared to when they were included in separate models. This suggests the interdependency among the brain, genes, social-demographic and psychological factors as proposed by the RDoC^6, 7^. Third, under no circumstances should the results of this paper be interpreted as entailing a value judgement about how people vary in measurements of cognitive abilities. Indeed, it is important to reflect on the fact that the way we measured cognitive abilities, e.g. through the *g*-factor here, reflects norms that are entrenched in cultures and societies of a certain time in history, rather than reflecting some universal truth or a supra-historical marker of cognitive abilities^5^. The value of the *g*-factor here is as a marker (present in early life) of a series of other important life outcomes in current societal circumstances.

In conclusion, we developed brain-based stacked, predictive models for children’s cognitive abilities that were longitudinally stable, generalisable and robust against missingness. More importantly, our brain-based models were able to partially mediate the relationships of childhood cognitive abilities with the socio-demographic, psychological and genetic factors. Accordingly, our approach should pave the way for future researchers to employ multimodal MRI as a useful research tool for integrative, RDoC-inspired research in cognition and mental health.

## Acknowledgments

Data used in the preparation of this article were obtained from the Adolescent Brain Cognitive Development (ABCD) Study (https://abcdstudy.org), held in the NIMH Data Archive (NDA). This is a multisite, longitudinal study designed to recruit more than 10,000 children age 9-10 and follow them over 10 years into early adulthood. The ABCD Study is supported by the National Institutes of Health and additional federal partners under award numbers U01DA041022, U01DA041028, U01DA041048, U01DA041089, U01DA041106, U01DA041117, U01DA041120, U01DA041134, U01DA041148, U01DA041156, U01DA041174, U24DA041123, U24DA041147, U01DA041093, and U01DA041025. A full list of supporters is available at https://abcdstudy.org/federal-partners.html. A listing of participating sites and a complete listing of the study investigators can be found at https://abcdstudy.org/scientists/workgroups/. ABCD consortium investigators designed and implemented the study and/or provided data but did not necessarily participate in the analysis or writing of this report. This manuscript reflects the views of the authors and may not reflect the opinions or views of the NIH or ABCD consortium investigators. We thank the developers of several R libraries, including semTools (Sunthud Pornprasertmanit), eNetXplorer (Julián Candia), and ggseg (Athanasia M. Mowinckel), for their technical advice. N.P and Y.W. were supported by Health Research Council Funding (21/618) and by University of Otago

## Supplementary Materials

### Appendix 1: g-Factor Based on Different Confirmatory Factor Analysis Models

Here using the first-layer training data, we compared factor scores of the g-factor across three different confirmatory factor analysis (CFA) models: the 2^nd^-order model, the single-factor model, and the mixture between Exploratory Factor Analysis (EFA) and CFA models, here by referred to as the EFA-CFA model.

For the 2^nd^-order model (see Supplementary Figure 1), we had the *g*-factor as the 2^nd^-order latent variable. We also had three 1^st^-order latent variables in the model: language (underlying Picture Vocabulary and Oral Reading Recognition), mental flexibility (underlying Flanker and Pattern Comparison Processing), and memory recall (underlying Picture Sequence Memory and Rey-Auditory Verbal Learning). This 2^nd^-order model of the *g*-factor showed a good fit: (a) scaled, robust Comparative Fit Index (CFI) =.995, (b) scaled, robust Tucker-Lewis Index (TLI) =.988, (c) scaled, robust Root Mean Square Error of Approximation (RMSEA) = .029 (90%CI=.015-.043) and (d) robust Standardized Root Mean Square Residual (SRMR) = .014. The *g*-factor latent variable of the 2^nd^-order model also had high internal consistency: OmegaL2=.78. The 2^nd^-order model led to the sample-size adjusted Bayesian (BIC) at 46549.957.

**Supplementary Figure 1.**
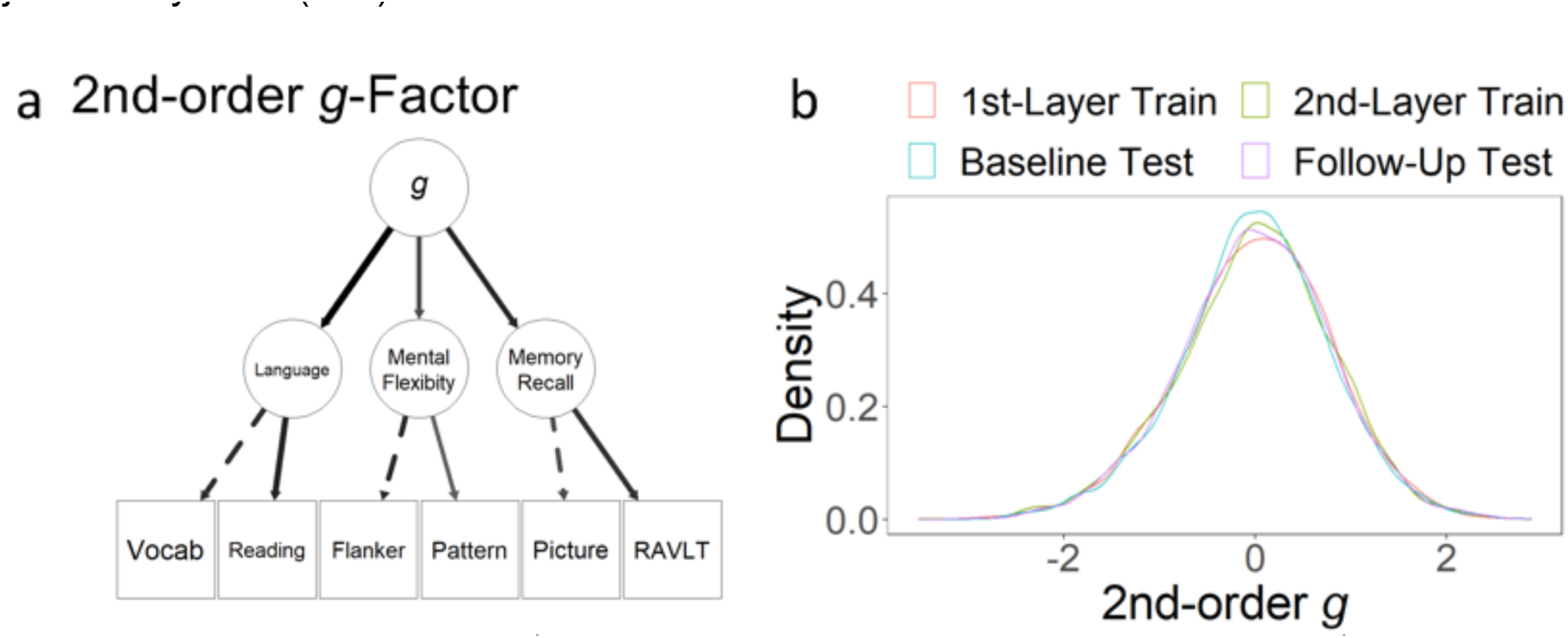
The 2^nd^-order model of the g-factor. 2A shows the 2^nd^-order model of the g-factor. Line thickness reflects the magnitude of standardized parameter estimates. The dotted lines indicate marker variables that were fixed to 1. Vocab = Picture Vocabulary; Reading = Oral Reading Recognition; Pattern = Pattern Comparison Processing; Picture = Picture Sequence Memory; RAVLT = Rey-Auditory Verbal Learning. 2B shows the distribution of the g-factor factor score across the four data splits. The distribution of the g-factor factor scores was similar across data splits.

For the single-factor model (see Supplementary Figure 2), we had the *g*-factor as the only one latent variable, underlying variation in all manifest variables: Picture Vocabulary, Oral Reading Recognition, Flanker and Pattern Comparison Processing, Picture Sequence Memory and Rey-Auditory Verbal Learning. This single-factor model of the *g*-factor showed a numerically poorer fit: (a) scaled, robust Comparative Fit Index (CFI) =.828, (b) scaled, robust Tucker-Lewis Index (TLI) =.713, (c) scaled, robust Root Mean Square Error of Approximation (RMSEA) = .139 (90%CI=.129-.149) and (d) robust Standardized Root Mean Square Residual (SRMR) = .070. The single-factor model led to a poorer (i.e., higher) sample-size adjusted Bayesian (BIC) than the 2^nd^-order model, at 47028.981.

**Supplementary Figure 2.**
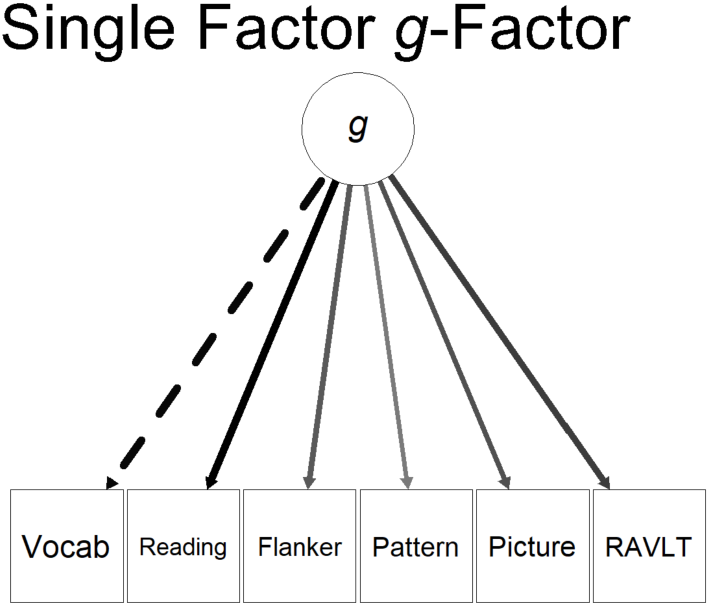
The single-factor model of the g-factor. Line thickness reflects the magnitude of standardized parameter estimates. The dotted lines indicate marker variables that were fixed to 1. Vocab = Picture Vocabulary; Reading = Oral Reading Recognition; Pattern = Pattern Comparison Processing; Picture = Picture Sequence Memory; RAVLT = Rey-Auditory Verbal Learning.

For the EFA-CFA model (see Supplementary Figure 3), we first applied EFA to the performance across the six cognitive tasks via the psych package. To implement EFA, we first ran a parallel analysis to determine the number of factors to retain (see Supplementary Figure 3a). The parallel analysis suggested three as the number of factors. We then ran EFA with three factors using ‘oblimin’ as the rotation, ‘maximum likelihood’ as the factoring method and “Thurstone” as the factor scoring method. This resulted in the same three factors used in the 2^nd^-order CFA model (see Supplementary Figure 3b). See Supplementary Table 1 for the standardized loadings (pattern matrix) based upon the correlation matrix. We then extracted factor scores from the final EFA model and used them as the manifest variables in another CFA model (see Supplementary Figure 3c).

Perhaps due to the use of EFA factor scores as the manifest variables, the EFA-CFA model resulted in a model with the perfect fit (as reported by the lavaan package): (a) scaled, robust Comparative Fit Index (CFI) = 1, (b) scaled, robust Tucker-Lewis Index (TLI) = 1, (c) scaled, robust Root Mean Square Error of Approximation (RMSEA) = 0 (90%CI= 0-0) and (d) robust Standardized Root Mean Square Residual (SRMR) = 0. The sample-size adjusted Bayesian (BIC) of the EFA-CFA was the best (i.e., lowest) among the three models at 17197.366.

**Supplementary Figure 3.**
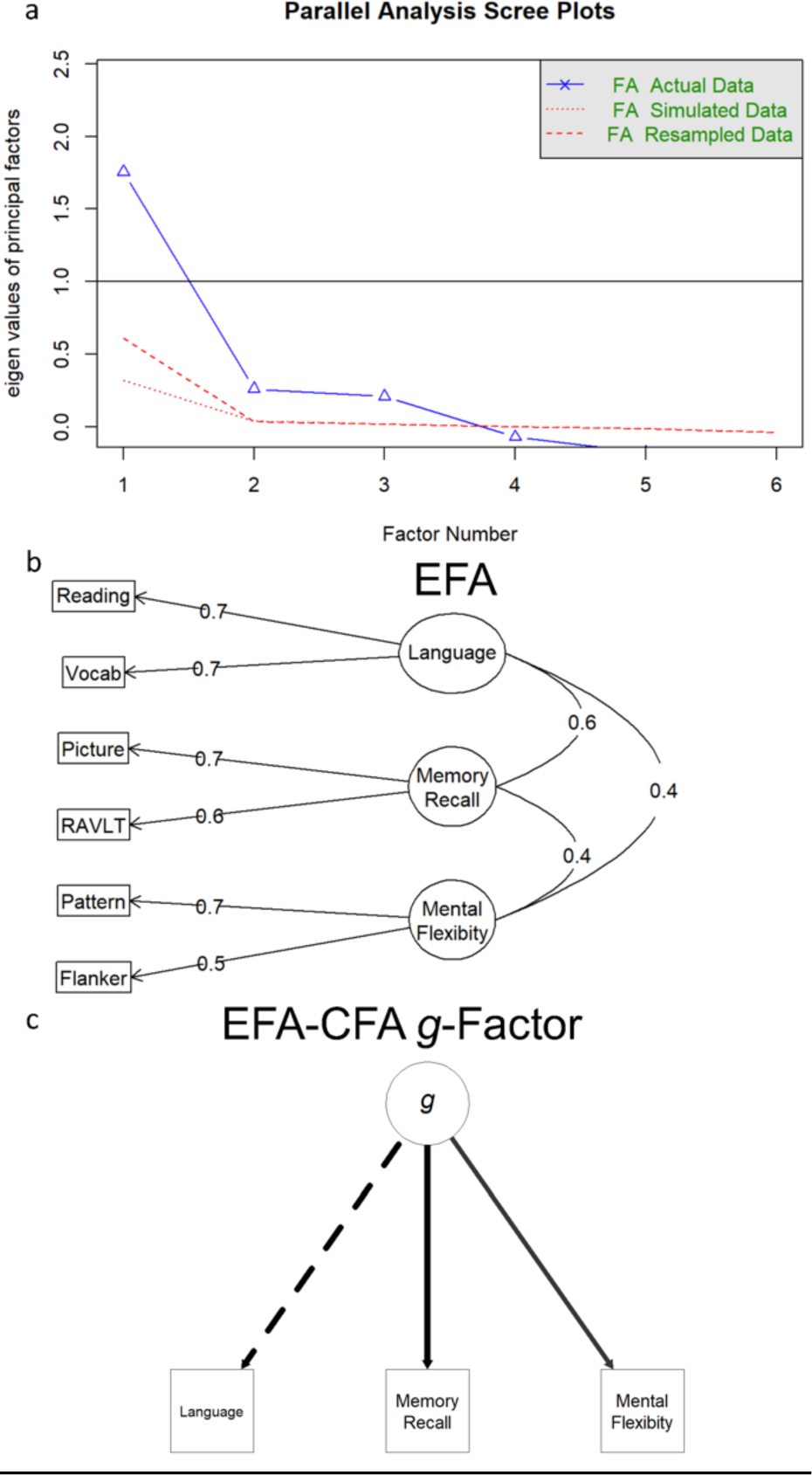
The EFA-CFA model of the g-factor. 3A shows the parallel analysis scree plot, suggesting three factors as the most likely solution for the EFA model. 3B shows the three-factor EFA model. 3C shows the EFA-CFA model, where its manifest variables were factor scores of the EFA model. Line thickness reflects the magnitude of standardized parameter estimates. The dotted lines indicate marker variables that were fixed to 1. Vocab = Picture Vocabulary; Reading = Oral Reading Recognition; Pattern = Pattern Comparison Processing; Picture = Picture Sequence Memory; RAVLT = Rey-Auditory Verbal Learning.

**Supplementary Table 1.**
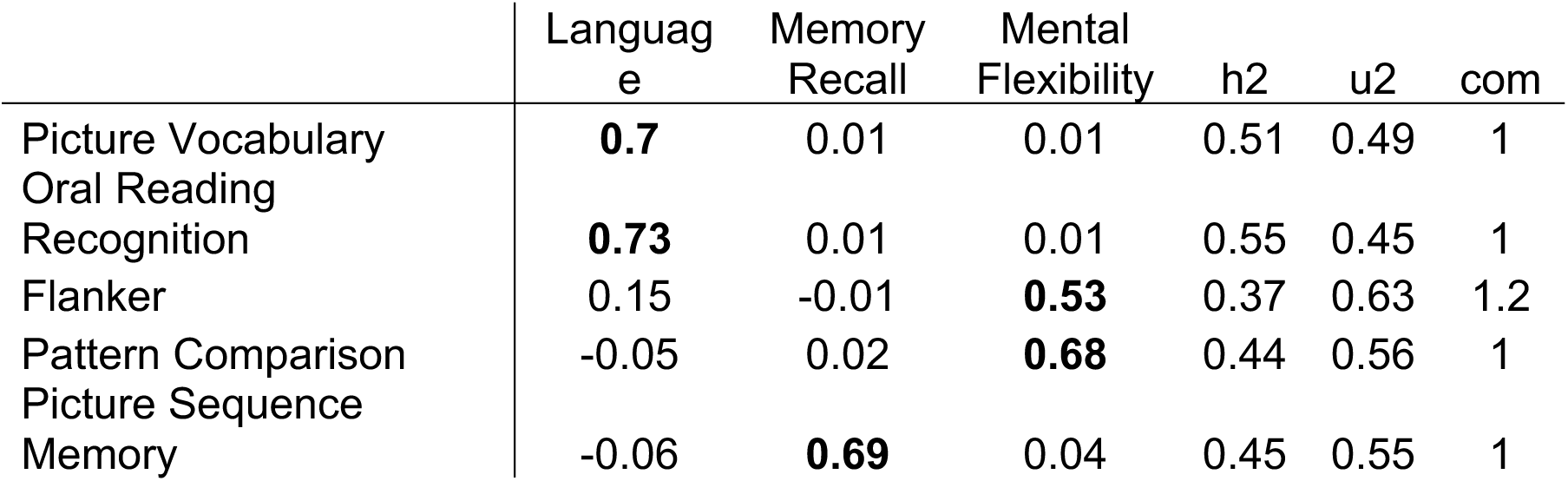
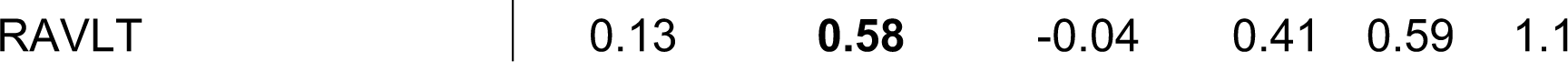
Standardized loadings based upon correlation matrix

We then examined the similarity in the factor scores of the *g*-factor based on three different CFA models at different data splits. We first extracted factor scores from the data used for fitting the CFA models, the 1^st^-layer training set, as well as unseen data sets, including the 2^nd^-layer training data, baseline test set and follow-up test data. Across data sets, we found high similarity in the factor scores of the *g*-factor across the three different CFA models at Pearson’s *rs* ≥ .987 (see Supplementary Figure 4). Accordingly, the choice of *g*-factor models had only minimal effects on the estimation of the factor scores for the *g*-factor.

**Supplementary Figure 4.**
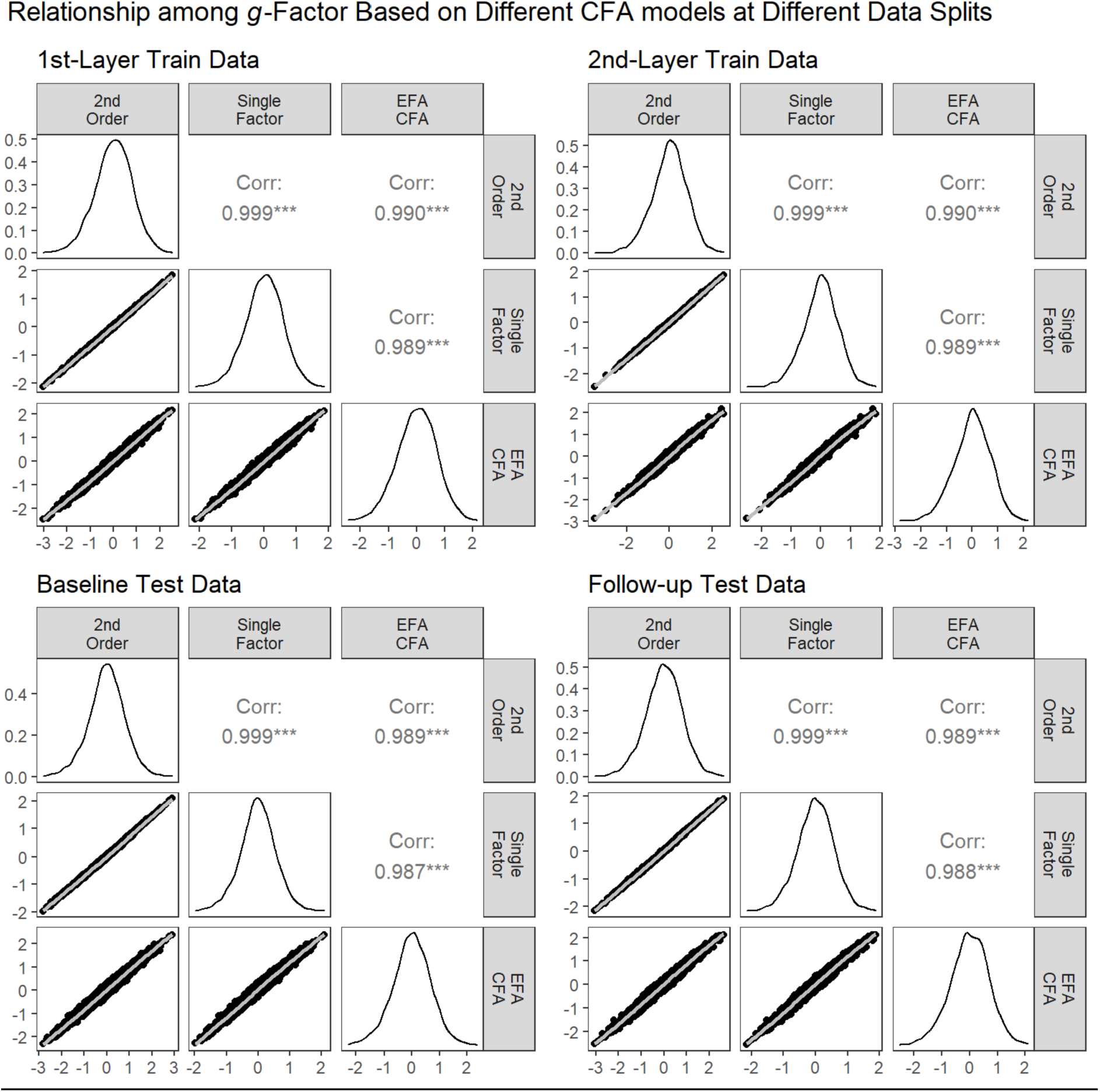
The Relationship in the factor scores of the g-factor based on three different CFA models at different data splits.

### Appendix 2: 2^nd^-order g-Factor based on different sets of data

To demonstrate the stability of the factor scores of the *g*-factor when applied to unseen data (i.e., not part of the modelling process), we compared the *g*-factor scores estimated from the first-layer training data vs. the scores estimated from the full baseline data. Using the first-layer training data to build the 2^nd^-order model led to highly similar factor scores of the *g*-factor to using the full baseline data across different data splits, at Pearson’s *rs* > .997 (see Supplementary Figure 5). Note we did not use the follow-up test data here since the follow-up test data included the same participants as the baseline test data, and including the data from the same participants twice may lead to a biased model.

**Supplementary Figure 5.**
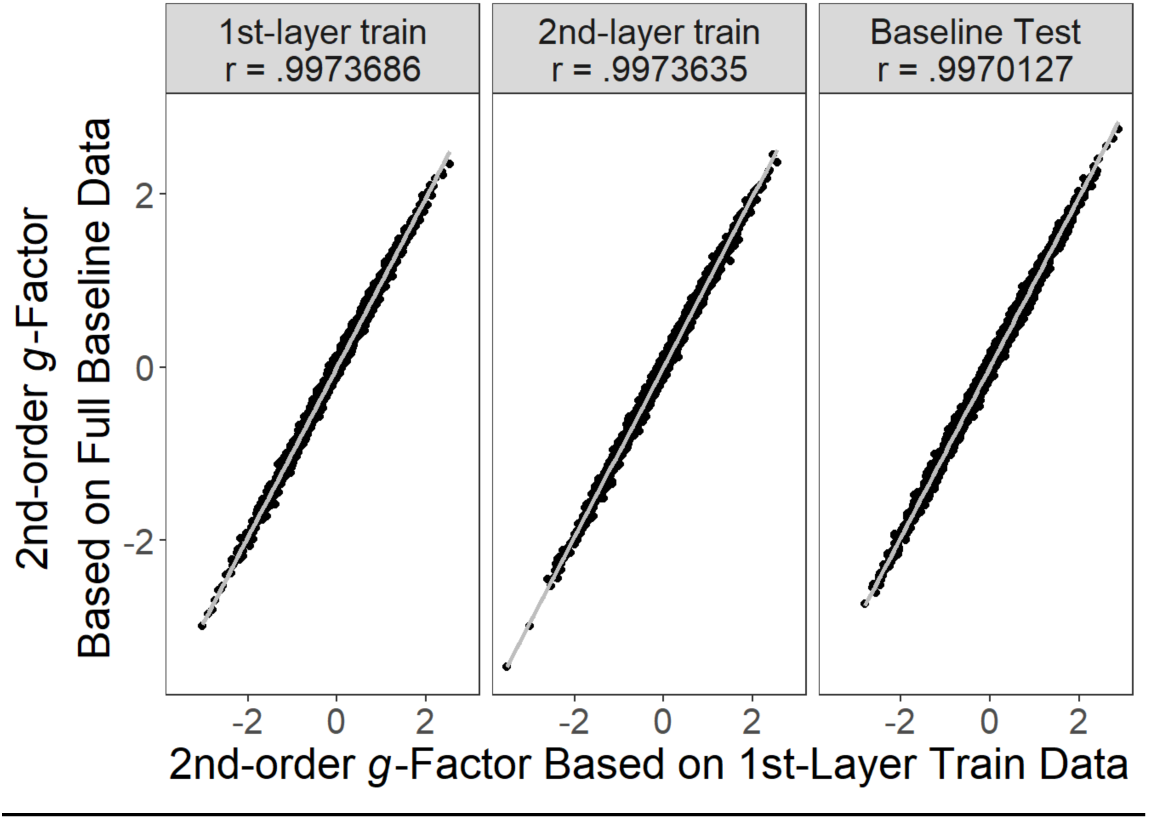
The similarity in the factor scores of the g-factor of the 2^nd^-order model based on whether the model was built from the 1^st^-layer training data or the full baseline data.

### Appendix 3: Hyperparameters for the Longitudinal Predictive Models for Multimodal MRI

From the first-layer training set, we found the best-tuned Elastic Net’s mixture was 0 for N-Back task-based fMRI, rs-fMRI and DIT, .1 for sMRI, and 1 for SST and MID task-based fMRI. Thus, whether to have brain features shrunk together (the Ridge solution, mixture close to 0) or to have certain features from all features selected (the Lasso solution, mixture close to 1) depended on the modality. The averaged Elastic Net’s penalty was .339 (SD = .45). From the second-layer training set, we found the best-tuned Random Forests’ mtry (the number of features randomly sampled at each path split) at 4 and min_n (the minimum number of observations in a node per path split) at 170.

## Notes

https://abcdstudy.org

### Competing Interest Statement

The authors have declared no competing interest.

